# Efficient replacement of long DNA fragments via non-homologous end joining at non-doding regions

**DOI:** 10.1101/2020.01.11.902791

**Authors:** Shan-Ye Gu, Jia Li, Jian-Bin Cao, Ji-Wen Bu, Yong-Gang Ren, Wen-Jie Du, Zhe-Cong Chen, Chu-Fan Xu, Min-Cang Wang, Lai Jiang, Cheng Huang, Jiu-Lin Du

## Abstract

Genomic DNA replacement for achieving sophisticated genetic manipulation is implemented currently through homogenous recombination/homology-dependent repair (HR/HDR). Here we report an efficient DNA fragment replacement method that is mediated by non-homologous end joining (NHEJ)-dependent DNA repair at two sites of CRISPR/Cas9-induced double-strand breaks at non-coding genomic regions flanking the exons of targeted genes. We demonstrated this method by generating three conditional alleles and two reporter lines of zebrafish. Functional assays of the conditional alleles proved that the genomic sequence between two inserted *loxP* sites was deleted by the Cre recombinase, and the phenotype after Cre-induced excision was comparable to previously reported mutants or morphants. Furthermore, combining double-fluorescence expression donor vectors, we showed that the efficiency of this NHEJ-mediated DNA replacement was around 3 times larger than that of HR/HDR-mediated approach. Our method provides a feasible strategy for genomic DNA replacement in zebrafish, which can be applicable for other organisms as well.

---

Genomic DNA replacement through homogenous recombination (HR) is a common approach in generating genetically engineered animals^1^. Due to its low efficiency, the HR-mediated approach usually requires both positive and negative selective systems to pre-screen gene-modified events in cultured embryonic stem cells^2^. In recent years, targeted double-strand breaks (DSBs) in genomes can be introduced efficiently by endonucleases^3-7^, including zinc-finger nucleases, transcription activator-like effector nucleases, and clustered regularly interspaced palindromic repeats (CRISPR)/Cas9. After DSBs, DNA repair is mainly via homology-directed repair (HDR) and/or non-homologous end joining (NHEJ)^8,9^. It was recently reported that genomic DNA replacement can be achieved via HDR at the site of DSBs in multiple organisms, including mouse and zebrafish^9-15^, but the efficiency is still not enough for general application, in particular for long DNA fragment replacement in zebrafish.

As NHEJ is 10-fold more active than HDR at DSB sites^9,16,17^, we speculated that NHEJ can be utilized to implement genomic DNA replacement with high efficiency. In this study, we developed a novel DNA replacement method that is mediated by NHEJ-dependent DNA repair at two sites of CRISPR/Cas9-induced DSBs at non-coding genomic regions flanking exons of the targeted gene. Using this methods, we generated conditional alleles in three zebrafish genomic loci (*th, kdrl* and *tcf3a*), which are known to be important for the development or function of the neural system, vascular system and muscle^18-20^. Then we confirmed the function of the conditional alleles by Cre-induced genomic sequence excision and phenotype characterization. Furthermore, we generated *th* and *gfap* zebrafish reporter lines, in which EGFP expression well recaptured the pattern of endogenous gene expression. Importantly, we compared the efficiency of NHEJ-mediated DNA replacement with that of HDR-mediated approach by using double-fluorescence expression donor vector. The efficiency of this NHEJ-mediated replacement of long DNA fragments is ∼ 3.7 - 5.3 times larger than those of HDR-mediated approaches. Taken together, we provide a novel and efficient strategy for achieving long genomic DNA replacement, making the sophisticated genome editing achievable to the zebrafish community.

## Results

### Generation of *th* conditional alleles via NHEJ-mediated genomic DNA replacement

To develop NHEJ-mediated DNA replacement method, we firstly tried to make a conditional allele of the zebrafish *tyrosine hydroxylase* (*th*) gene, which is specifically expressed in both dopaminergic neurons and noradrenergic neurons. Two single guide RNAs (sgRNAs) were designed to specifically target two different non-coding sites in the 7^th^ (for sgRNA1) and 8^th^ (for sgRNA2) introns of the *th* with a cleavage efficiency of 59% and 68%, respectively (Fig. 1a and Supplementary Fig. 1a). We constructed a *th* conditional donor (*th-loxP-exon8-loxP*) consisting of a left arm, a *loxP-exon8-loxP* fragment, and a right arm (middle in Fig. 1a). The left arm includes the upstream sequence of the 5’ side of the sgRNA1 target in the 7^th^ intron and the sequence of the sgRNA1 target, the right arm includes the sequence of the sgRNA2 target and the downstream sequence of the 3’ side of the sgRNA2 target in the 8^th^ intron, and the *loxP-exon8-loxP* fragment contains the sequence between the two sgRNA targets in the *th* locus and the sequence of two *loxP* sites flanking the 8^th^ exon. We then co-injected the donor with zebrafish optimized Cas9 mRNA (zCas9 mRNA) and the two sgRNAs into one-cell-stage embryos of the knockin zebrafish line Ki(th-P2A-EGFP), in which the *P2A* and *enhanced green fluorescence protein* (*EGFP*) are sequentially ligated to the 3’ side of the *th* coding sequence and thus th-expressing cells are labeled by EGFP^21^, and raised them to adulthood for screening the founders. Successful DNA replacement in the *th* locus with the *loxP-exon8-loxP* fragment in progenies of F0 was verified by PCR using *th* locus-specific (F1 and R1) and donor-specific (F2 and R2) primers (Supplementary Fig. 1b), and sequencing of PCR products amplified across the entire edited region with *th* locus-specific primers (Fig. 1b and Supplementary Fig. 1c). In a total of 41 F0 screened, we identified 3 replacement founders [Ki(th^fl^-P2A-EGFP)] with an 18% ± 9% F1 progeny carrying the replacement event, which was determined by PCR of F1 progenies one by one (Table 1, Supplementary Fig. 2 and Supplementary Table 1). The indels at both the 5’ and 3’ junctions indicate that the DNA replacement occurred is through NHEJ (Fig. 1b, Table 1 and Supplementary Fig. 1c).

**Table 1.**
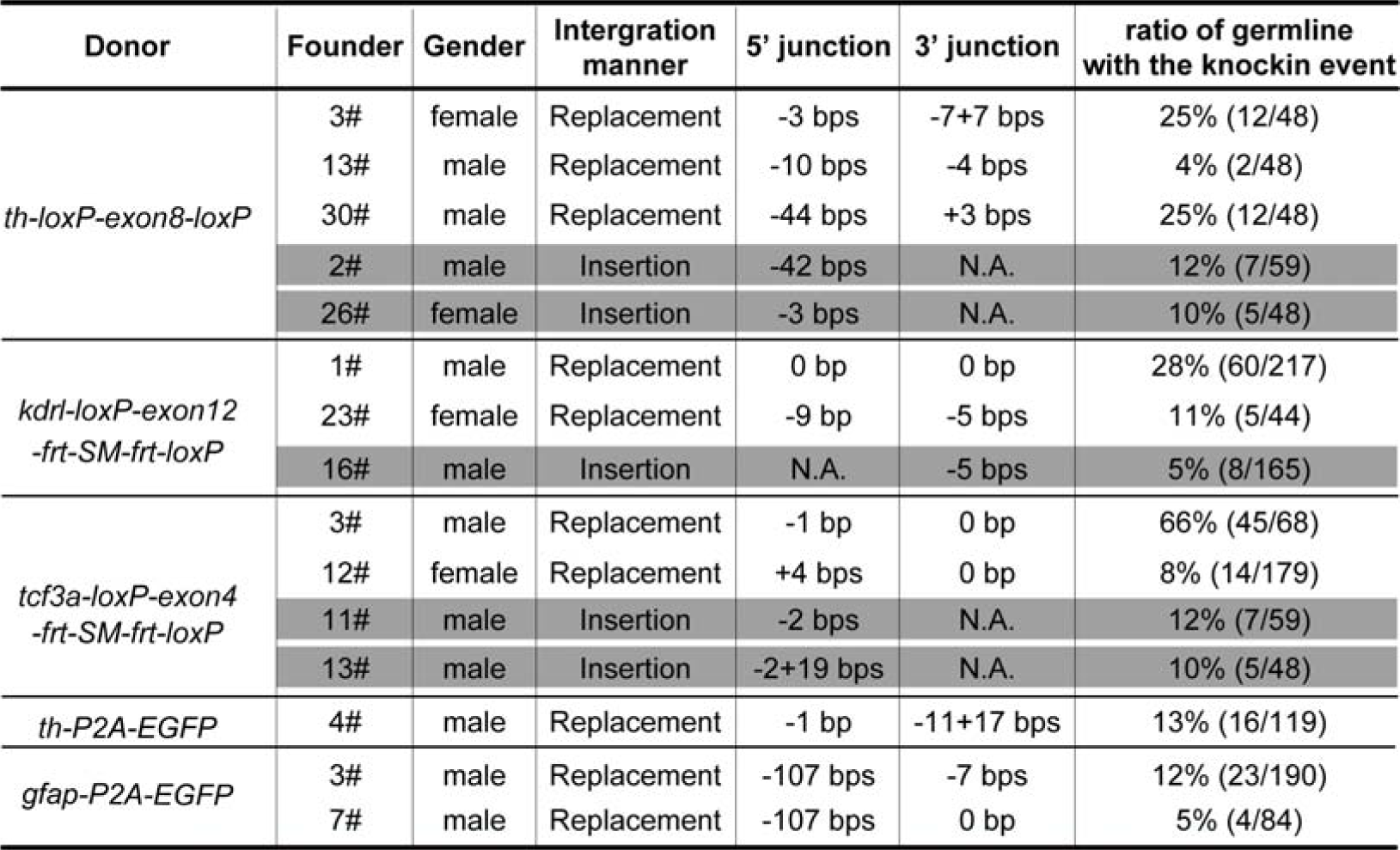
Information of edited alleles and germline mosaicism ratio of the founders.

**Fig. 1.**
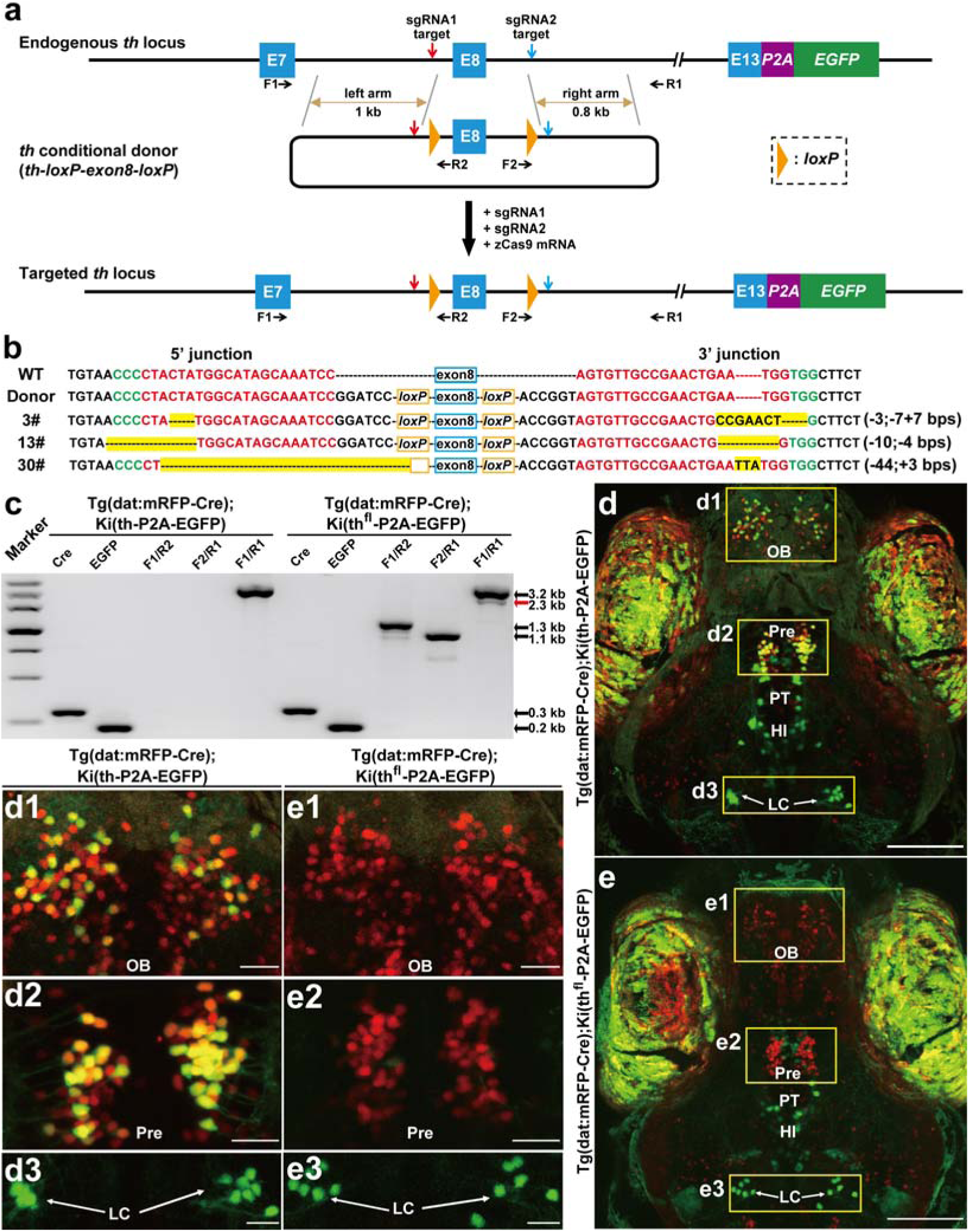
Generation of zebrafish *th* conditional alleles via NHEJ-mediated DNA replacement. **a** Schematic of the strategy for NHEJ-mediated DNA replacement at the *th* locus of Ki(th-P2A-EGFP) zebrafish, in which the coding sequences of *P2A* and *EGFP* are sequentially ligated to the 3’ side of the *th* coding sequence and thus EGFP is expressed in th-expressing cells. The sgRNA1 and sgRNA2 targets are indicated by red or blue arrows, respectively. In the *th* conditional donor (*th-loxP-exon8-loxP*), two *loxP* sites are added and homology arms are indicated by brown lines with double arrows. The length of the left and right arms is 999 base pairs (bps) and 831 bps, respectively. The zebrafish *th* has 13 exons, and E7, E8 and E13 represent the 7^th^, 8^th^ and 13^th^ exons, respectively. The length of E8 is 136 bps. The endogenous E8 is replaced by the *th-loxP-exon8-loxP* cassette in the *th* locus after co-injecting the donor with the sgRNA1, sgRNA2 and zCas9 mRNA. The primers (F1, R1, F2, R2) for verification are indicated by short black arrows. **b** 5′ and 3′ junction sequences of F1 progenies of three founders carrying DNA replacement (3#, 13# and 30#) in comparison with the sequences of the wild-type (WT) animal and the donor vector. The indel mutations are highlighted in yellow, and the PAM and sgRNA target sequences are shown in green and red, respectively. The numbers of insertion (+) and/or deletion (-) of base pairs at the 5’ and 3’ junctions are shown in the brackets at right. **c** The entire edited region PCR verification of conditional knockout, showing an additional product at ∼ 2.3 kb (red arrow) amplified by the primers F1 and R1 in heterozygous [Tg(dat:mRFP-Cre);Ki(th^fl^-P2A-EGFP)] larvae. This product was in principle excised from the original 3.2 kb fragment by Cre (see **a**). **d, e** Projected *in vivo* confocal images (dorsal view) of [Tg(dat:mRFP-Cre);Ki(th-P2A-EGFP)] (**d**) and [Tg(dat:mRFP-Cre);Ki(th^fl^-P2A-EGFP)] (**e**) larvae at 5 days post-fertilization (dpf), showing that in the conditional knockout embryo, mRFP-Cre-expressing cells (in red) do not express EGFP. In **e**, the larva was the progeny of the founder 3#. The enlarged views of the areas outlined by the rectangles in **d** and **e** are shown in **d1-d3** and **e1-e3**, respectively. To clearly visualize each nucleus, the range of z-axis for projection is different among images. In comparison with **d** and **e**, the projected images in **d1** and **e1** did not include the 12 most dorsal layers which contain non-specific signals on the skin, and the projected images in **d2** and **e2** did not include the 18 most ventral layers where cells do not express mRFP-Cre. HI, intermediate hypothalamus; LC, locus coeruleus; OB, olfactory bulb; Pre, pretectum; PT, posterior tubercular. Scale bars: 100 µm (**d, e**), 20 µm (**d1-d3, e1-e3**).

### Functional validation of *th* conditional alleles

To examine the excision by Cre, we did PCR with F1 and R1 in larvae generated by crossing conditional allele Ki(th^fl^-P2A-EGFP) with Tg(dat:mRFP-Cre), in which Cre fused to the C terminal of the monomer red fluorescent protein (mRFP) is specifically expressed in dopaminergic neurons via the promoter of *dopamine transporter* (*dat*). After checking the PCR products of across the entire edited region, we found there was a band with ∼ 0.9 kilo base pairs (bps) smaller than control (red arrow in Fig. 1c). This reduction size was equivalent to the length of the fragment between two *loxP* sites (823 bps), suggesting correct excision by Cre.

To further test the functionality of the targeted genomic locus, we first examined the EGFP expression pattern in the larvae of Ki(th^fl^-P2A-EGFP) and found that it was similar to Ki(th-P2A-EGFP), indicating that NHEJ-mediated replacement does not affect the integrity of the *th-P2A-EGFP* (Supplementary Fig. 3). Then we used Cre recombinase to examine whether the conditional allele generated through NHEJ-mediated DNA replacement works. As a control, we crossed Ki(th-P2A-EGFP) with Tg(dat:mRFP-Cre). As expected, EGFP and mRFP were co-expressed in some dopaminergic neurons in the olfactory bulb (OB) and pretectum (Pre) in double heterozygote progenies (Fig. 1d, 1d1 and 1d2). However, in embryos of Ki(th^fl^-P2A-EGFP) zebrafish crossed with Tg(dat:mRFP-Cre), EGFP expression was not observed in mRFP-positive cells in the Pre and OB (Fig. 1e, 1e1 and 1e2), confirming the occurrence of the frame-shift mutation induced by excising the 8^th^ exon (136 bps in length) between the two *loxP* sites in the *th* locus. As a control, EGFP expression was not affected in noradrenergic neurons in the locus coeruleus (LC) where mRFP-Cre did not express (Fig. 1d, 1d3, 1e and 1e3). Taken together, these results indicate that the conditional allele generated by NHEJ-mediated DNA replacement is functional.

### Generation of *kdrl* conditional alleles via NHEJ-mediated genomic DNA replacement

To demonstrate the general applicability of this NHEJ-mediated DNA replacement, we next tried the zebrafish *kinase insert domain receptor like* (*kdrl*) gene, which is specifically expressed in vascular endothelial cells (ECs). Two sgRNA flanking the 12^th^ exon of *kdrl* were selected with a cleavage efficiency about 41% and 71%, respectively (Fig. 2a and Supplementary Fig. 4a). To reduce the labor of intensive screening of conditional alleles, we introduced a 1.2-kb selective marker (SM) in the designed replacement fragment. The SM sequentially consisted of the cardiac *myl7* promoter, coding sequence of DsRed and SV40 polyA signal, and was flanked by *frt* recombination sites. Thus *DsRed* can be specifically translated in cardiomyocytes and excised by Flp recombinase. We constructed a SM-containing *kdrl* conditional donor (*kdrl-loxP-exon12-frt-SM-frt-loxP*, middle in Fig. 2a), in which the transcriptional direction of SM was contrary to the endogenous *kdrl* gene to permit *kdrl* transcription normally. We co-injected zCas9 mRNA, the two sgRNAs and the donor into one-cell-stage embryos of Tg(kdrl:EGFP) transgenic zebrafish, in which vascular ECs were labeled by EGFP, and raised the embryos with broad expression of DsRed in the heart to adulthood (Supplementary Fig. 4b). We identified 2 founders [Ki(kdrl^fl^)] out of 25 F0 screened that carried DNA replacement, as verified by PCR of the entire edited region and sequencing (Table 1, Supplementary Fig. 5a, b and Supplementary Table 1). To confirm that the SM can be removed, we injected Flp mRNA into one-cell-stage embryos generated via Ki(kdrl^fl/+^) intercross and found that 43 out of 47 injected larvae at 3 days post-fertilization (dpf) did not express DsRed in the heart, indicating the occurrence of Flp-induced SM excision. The Flp-induced excision of SM was also confirmed by PCR of the entire edited region and sequencing (Fig. 2c and Supplementary Fig. 5c).

**Fig. 2.**
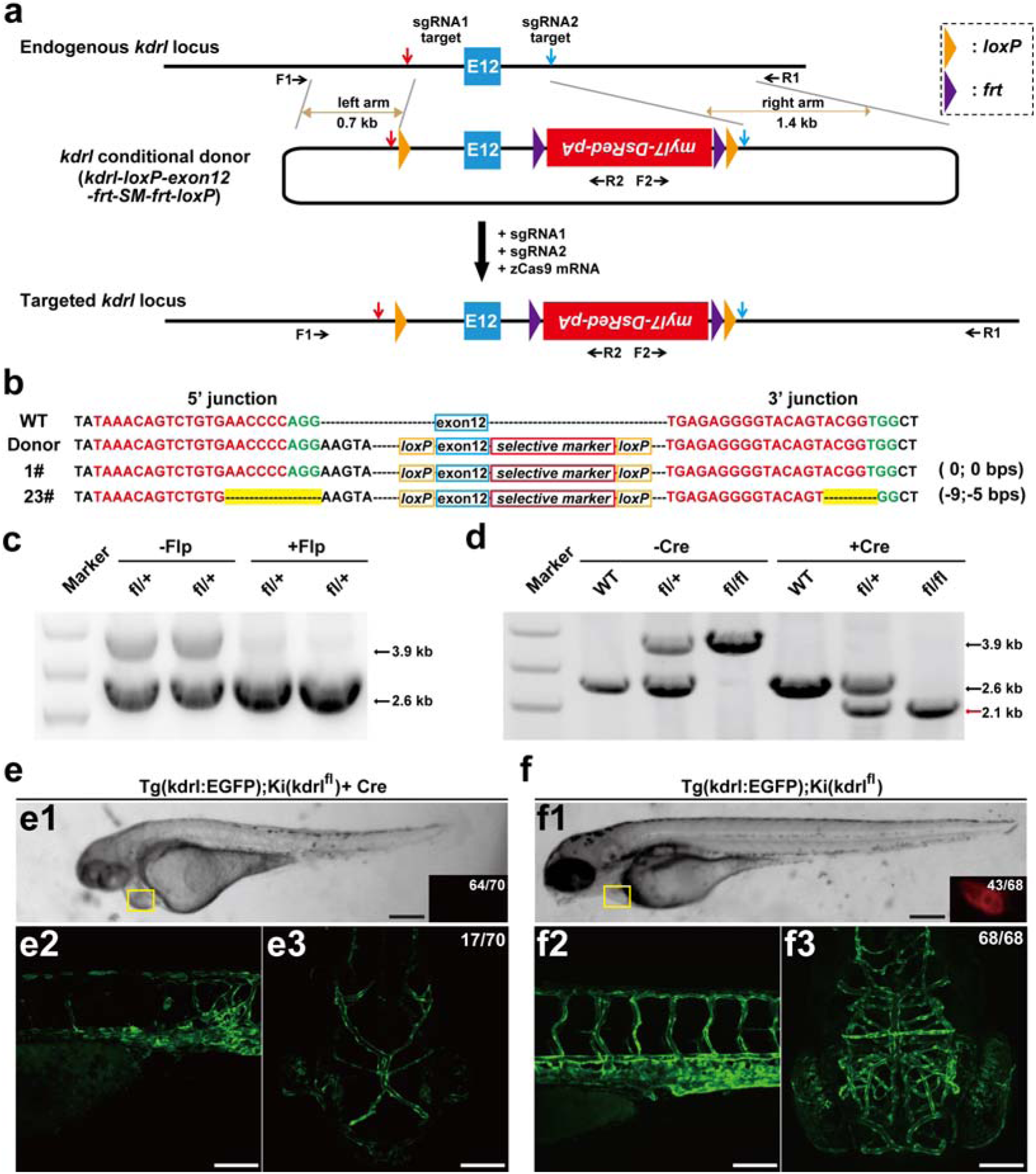
Generation of zebrafish *kdrl* conditional alleles with a selective marker via NHEJ-mediated DNA replacement. **a** Schematic of the strategy for NHEJ-mediated DNA replacement at the *kdrl* locus of Tg(kdrl:EGFP) zebrafish, in which the vascular endothelial cells express EGFP. The sgRNA1 and sgRNA2 targets are indicated by red or blue arrows, respectively. In the *kdrl* conditional donor (*kdrl-loxP-exon12-frt-SM-frt-loxP*), two loxP sites and homology arms (brown lines with double arrows) were added. The length of the left and right arms is 686 bps and 1297 bps, respectively. A selective marker (SM) can expresses DsRed under the control of the *myl7* promoter, and its transcriptional direction is contrary to *kdrl* transcriptional direction. The zebrafish *kdrl* has 30 exons, and E12 (12^th^ exon) contains 109 bps. The endogenous E12 is replaced by the *kdrl-loxP-exon12-frt-SM-frt-loxP* cassette at the *kdrl* locus after co-injecting the donor with the sgRNA1, sgRNA2 and zCas9 mRNA. The primers (F1, R1, F2, R2) for verification are indicated by short black arrows. **b** 5′ and 3′ junction sequences of F1 progenies of the two Ki(kdrl^fl^) founders in comparison with the sequences of WT zebrafish and the donor vector. The indel mutations are highlighted in yellow, and the PAM and sgRNA target sequences are shown in green and red, respectively. The numbers of insertion (+) and/or deletion (-) of bps at the 5’ and 3’ junctions are shown in the brackets at the right. **c** The entire edited region PCR analysis for the SM deletion in 3-dpf heterozygote Ki(kdrl^fl^) larvae with Flp mRNA injection at one-cell stage. **d** The entire edited region PCR analysis of heterozygote Ki(kdrl^fl^) intercross larvae without (“-Cre”) or with (“+Cre”) Cre mRNA injection at one-cell stage. The red arrow indicates the additional product at ∼ 2.1 kb, which were in principle excised from the original ∼ 3.9 kb fragment by Cre. **e, f** *In vivo* bright-filed (top) and projected confocal (bottom) images of 3-dpf [Tg(kdrl:EGFP);Ki(kdrl^fl/+^)] intercross larvae with (**e**) or without (**f**) Cre mRNA injection at one-cell stage, showing that Cre mRNA injection causes the loss of DsRed expression in the heart (inset in **e1**) and the defects of blood vessels in the trunk (**e2**) and brain (**e3**). Scale bars: 250 µm (**e1, f1**), 100 µm (**e2, e3, f2, f3**).

### Functional validation of *kdrl* conditional alleles

To test the functionality of the *kdrl* conditional allele, we injected Cre mRNA at one-cell-stage embryos of [Tg(kdrl:EGFP);Ki(kdrl^fl/+^)] intercross and then did PCR of the entire edited region and sequencing of individual larvae. The excision of the *kdrl* 12^th^ exon was observed (Fig. 2d and Supplementary Fig. 5d). As expected, the majority (64/70) of Cre-injected larvae did not express DsRed in the heart. Furthermore, we found that about one fourth (17/70) of Cre-injected larvae showed vascular defects in both the trunk and brain (Fig. 2e), consistent with the vascular phenotype of the *kdrl* mutant homozygote^19^. As a control, all (68/68) of [Tg(kdrl:EGFP);Ki(kdrl^fl/+^)] intercross larvae without Cre injection displayed normal vascular patterns (Fig. 2f). These results show that conditional alleles containing a selective marker is also achievable via NHEJ-mediated DNA replacement.

### Generation of *tcf3a* conditional alleles via NHEJ-mediated genomic DNA replacement

We further generated another SM-containing conditional allele of *tcf3a* (also named *E12*), which plays an important role in muscle development at early developmental stages in zebrafish^20^. Similar to the design of the *kdrl* conditional allele, we chose two sgRNA targets flanking the 4^th^ exon with a cleavage efficiency of 89% and 56%, respectively (Fig. 3a and Supplementary Fig. 6a). We got 2 replacement founders [Ki(tcf3a^fl^)] from 15 F0, as verified by PCR of the entire edited region and sequencing (Fig. 3b, Table 1, Supplementary Fig. 6b, c and Supplementary Table 1). Then we validated its functionality by Cre mRNA injection. The DNA sequence in the conditional allele of *tcf3a* was correctly excised (Fig. 3c) and the phenotype of larvae carrying Cre-induced *tcf3a* conditional knockout was comparable to *tcf3a* morphants (Fig. 3d and 3e; see also Ref. 20).

**Fig. 3.**
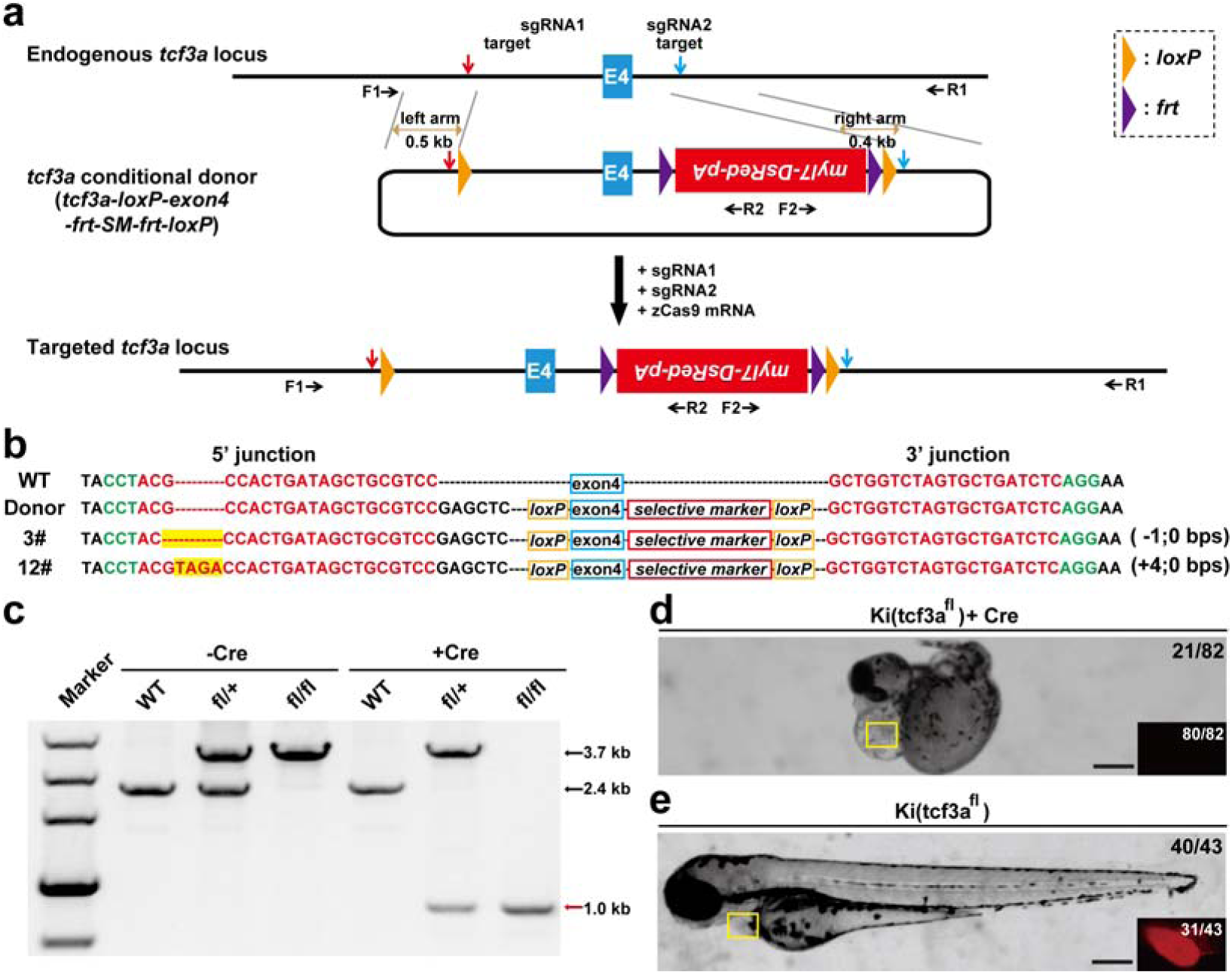
Generation of zebrafish *tcf3a* conditional alleles with a selective marker via NHEJ-mediated DNA replacement. **a** Schematic of the strategy for NHEJ-mediated DNA replacement at the *tcf3a* locus. The sgRNA1 and sgRNA2 targets are indicated by red or blue arrows, respectively. In the *tcf3a* conditional donor (*tcf3a-loxP-exon12-frt-SM-frt-loxP*), the length of the left and right arms is 530 bps and 424 bps, respectively. The length of E4 is 68 bps. **b** 5′ and 3′ junction sequences of F1 progenies of the two Ki(tcf3a^fl^) founders in comparison with the sequences of WT zebrafish and the donor vector. **c** The entire edited region PCR analysis of heterozygote Ki(tcf3a^fl^) intercross larvae without (“-Cre”) or with (“+Cre”) Cre mRNA injection at one-cell stage. The red arrow indicates the additional product at ∼ 1.0 kb, which were in principle excised from the original ∼ 3.7 kb fragment by Cre. **d, e** Bright-filed and fluorescent images of 2.5-dpf Ki(tcf3a^fl/+^) intercross larvae with (**d**) or without (**e**) Cre mRNA injection at one-cell stage, showing that Cre mRNA injection causes no expression of DsRed in the heart (inset in **d**) and the defects of gross morphology (**d**). Scale bars: 250 µm (**d, e**).

### Generation of a *th* reporter line via NHEJ-mediated genomic DNA replacement

To demonstrate that this NHEJ-mediated DNA replacement method can be used for making reporter lines, two sgRNAs were designed to specifically target other two sites in the *th* locus, one lying in the last intron (sgRNA3, see also Ref. 21) and the other overlapping the stop codon (sgRNA4) with a cleavage efficiency of 83% and 36%, respectively (Fig. 4a and Supplementary Fig. 7a). We constructed a *th* reporter donor (*th-P2A-EGFP*) that consists of a left arm, *P2A-EGFP* coding sequence, and a right arm (middle in Fig. 4a). As the sgRNA4 target partially overlaps with the 3’ side of the *th* coding sequence (CDS), we introduced silent mutations at the 3’ side of the *th* CDS in the *th-P2A-EGFP* donor vector (Supplementary Fig. 7b). Therefore, the 3’ CDS of the *th* in the donor cannot be excised by CRISPR/Cas9. We co-injected zCas9 mRNA, the two sgRNAs and donor vector into one-cell-stage wild-type embryos. We raised the injected embryos to adulthood and got 1 founder [Ki(th^EGFP^)] out of 11 F0 screened that carried NHEJ-mediated DNA replacement, as verified by PCR of the entire edited region, sequencing and EGFP expression pattern (Fig. 4b, c and Supplementary Fig. 7c, d).

**Fig. 4.**
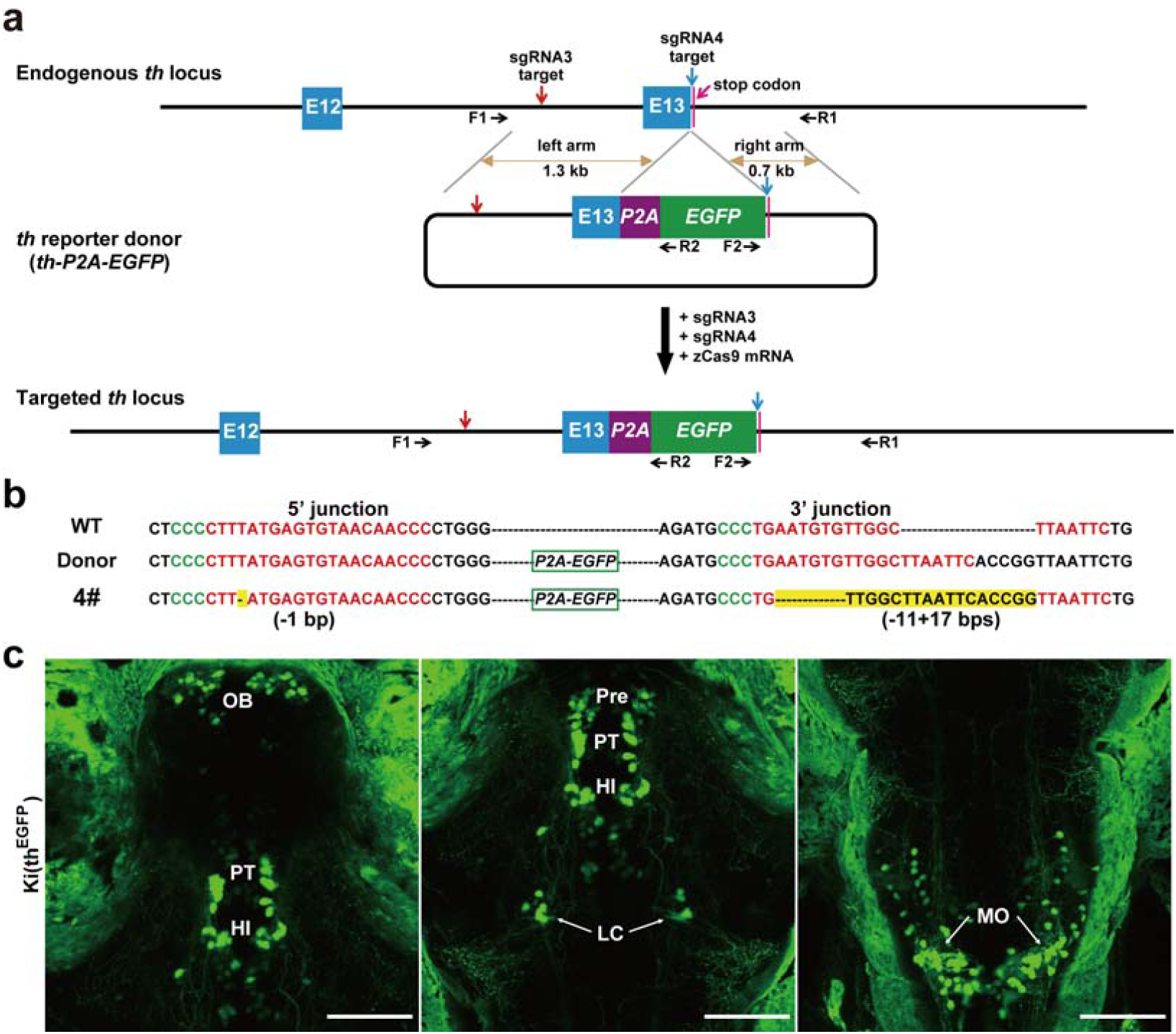
Generation of zebrafish *th* reporter line via NHEJ-mediated DNA replacement. **a** Schematic of NHEJ-mediated DNA replacement at the zebrafish *th* locus for generating EGFP reporter lines. The target sites of sgRNA3 and sgRNA4 are indicated by red or blue arrows, respectively. The *th* stop codon is indicated by a pink line. In the *th* reporter donor (*th-P2A-EGFP*), the homology arms are indicated by brown lines with double arrows. The length of the left and right arms is 1282 bps and 671 bps, respectively. The endogenous E13 in the *th* locus was replaced by the *th-P2A-EGFP* cassette after co-injection of the donor with the sgRNA3, sgRNA4 and zCas9 mRNA. The primers (F1, R1, F2, R2) for verification are indicated by short black arrows. **b** 5′ and 3′ junction sequences of F1 progeny of the Ki(th^EGFP^) founder in comparison with the sequence of WT zebrafish and the donor vector. The indel mutations are highlighted in yellow, and the PAM and sgRNA target sequences are shown in green and red, respectively. The numbers of insertion (+) and/or deletion (-) of bps at the 5’ and 3’ junctions are shown in the brackets. **c** Representative projected *in vivo* confocal images (dorsal view) of a 3-dpf Ki(th^EGFP^) larva at different visual fields. HI, intermediate hypothalamus; LC, locus coeruleus; MO, medulla oblongata; OB, olfactory bulb; Pre, pretectum; PT, posterior tubercular. Scale bars, 100 µm.

### Generation of a *gfap* reporter line via NHEJ-mediated genomic DNA replacement

We then demonstrated the applicability of this method by making another reporter line of the zebrafish *glial acidic fibrillary protein (gfap)* gene, which is specifically expressed in glial cells of the nervous system^21,22^. Two sgRNAs were selected to specifically target two different sites in the *gfap*, one in the last intron (sgRNA1) and the other in 3’ UTR region (sgRNA2) with a cleavage efficiency of 74% and 74%, respectively (Fig. 5a and Supplementary Fig. 8a). We co-injected the *gfap-P2A-EGFP* donor with zCas9 mRNA and the two sgRNAs, and observed EGFP expression in the spinal cord of ∼ 88% (160/181) larvae (Supplementary Fig. 8b and Supplementary Table 2). We raised 8 larvae with broad EGFP expression to adulthood and found 2 founders [Ki(gfap^EGFP^)] carried NHEJ-mediated DNA replacement, as verified by PCR of the entire edited region, sequencing and EGFP expression pattern (Fig. 5b, c, Supplementary Fig. 9 and Supplementary Table 1). These results indicate that NHEJ-mediated replacement can be readily applied for making reporter zebrafish lines.

**Fig. 5.**
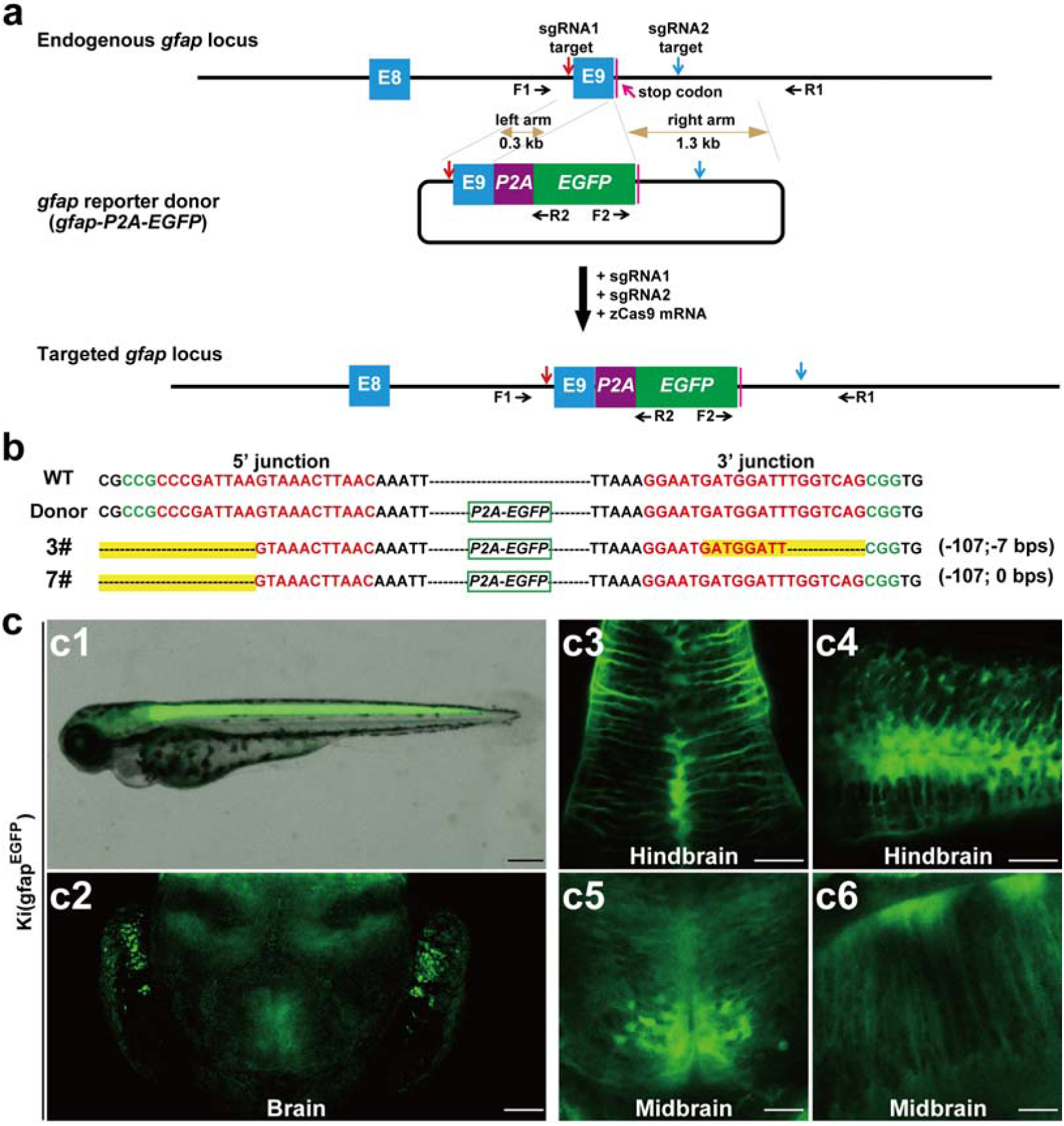
Generation of zebrafish *gfap* reporter line via NHEJ-mediated DNA replacement. **a** Schematic of the strategy for NHEJ-mediated DNA replacement at the zebrafish *gfap* locus. The *gfap* stop codon is indicated by a pink line. The homology arms are indicated by brown lines with double arrows, and the lengths of the left and right homology arms are 324 bps and 1326 bps, respectively. **b** 5′ and 3′ junction sequences of F1 progenies of the two Ki(gfap^EGFP^) founders in comparison with the sequence of WT zebrafish and the donor vector. **c** Representative projected *in vivo* confocal images of a 3-dpf Ki(gfap^EGFP^) larvae at different arears. Lateral view: **c1, c4, c6**; dorsal view: **c2, c3, c5**. Scale bars: 250 µm (**c1**), 50 µm (**c2**), and 25 µm (**c3-c6**).

### Comparison of the efficiency of genomic DNA replacement between NHEJ- and HDR-mediated approaches

To compare the efficiency of NHEJ- and HDR-mediated DNA replacement, we modified the *gfap-P2A-EGFP* donor by adding a negative SM (*gfap-P2A-EGFP-IRES-tdT*; Fig. 6a). The negative SM consisted of an internal ribozyme entry site (IRES), tdTomato (tdT) CDS and SV40 polyA signal (*IRES-tdT*), and was then linked to the 3’ side of a shortened right arm. In this right arm, the *gfap* transcriptional terminate signal was deleted to permit *IRES-tdT* transcription, and the target site of sgRNA2 was kept. We validated the *gfap-P2A-EGFP-IRES-tdT* donor vector by co-injecting with sgRNA1 and zCas9, and found that most of EGFP-expressing glial cells (197/219 from 16 larvae) also expressed tdT, indicating the insertion of the full-length of the donor vector at the sgRNA1 target site (Fig. 6b; see also Ref. 21). However, when co-injecting the donor vector with the two sgRNAs, the majority of EGFP-expressing glial cells did not express tdT (Fig. 6c), implying the occurrence of *P2A-EGFP* but not *IRES-tdT* integration in the *gfap* locus. To compare the difference in the efficiency of NHEJ- and HDR-mediated DNA replacement, we also constructed two HDR-mediated donors in which both the sgRNA1 and sgRNA2 targets were mutated at protospacer adjacent motifs (Fig. 6d, e, left). One HDR-mediated donor had same homology arms with the NHEJ-mediated donor (Fig. 6c, d), the other had both longer left and right homology arms (Fig. 6c, e). After co-injecting the NHEJ- or HDR-mediated *gfap* donor vector with the two sgRNAs, we calculated the ratio of the larvae containing EGFP-only-expressing glial cells and the average number of EGFP-only-expressing glial cells in each embryo. We found that both the ratios in embryos injected with the NHEJ donor were > 5.5 and 8.0 times as large as those with HDR donors with long homology arms, respectively (Fig. 6f, g). EGFP-only expression in HDR donor-injected embryos is in principle due to HDR-mediated DNA replacement. As multiple-fragment integration rarely occurs^23^, EGFP-only expression in NHEJ donor-injected embryos can be caused mainly through two possibilities. The first one is through expected NHEJ-mediated DNA replacement, and the other is due to NHEJ-mediated insertion of the *P2A-EGFP* fragment at the sgRNA1 target site. Considering the fact that we identified 10 founders carrying NHEJ-mediated DNA replacement and 5 founders carrying insertion during all of our experiments (Table 1), we roughly estimated that the NHEJ-mediated DNA replacement accounts for ∼ 67% of EGFP-only-expression events. Therefore, we reasoned that the efficiency of NHEJ-mediated DNA replacement in the *gfap* locus is ∼ 3.7 - 5.3 times as much as that of HDR-mediated DNA replacement.

**Fig. 6.**
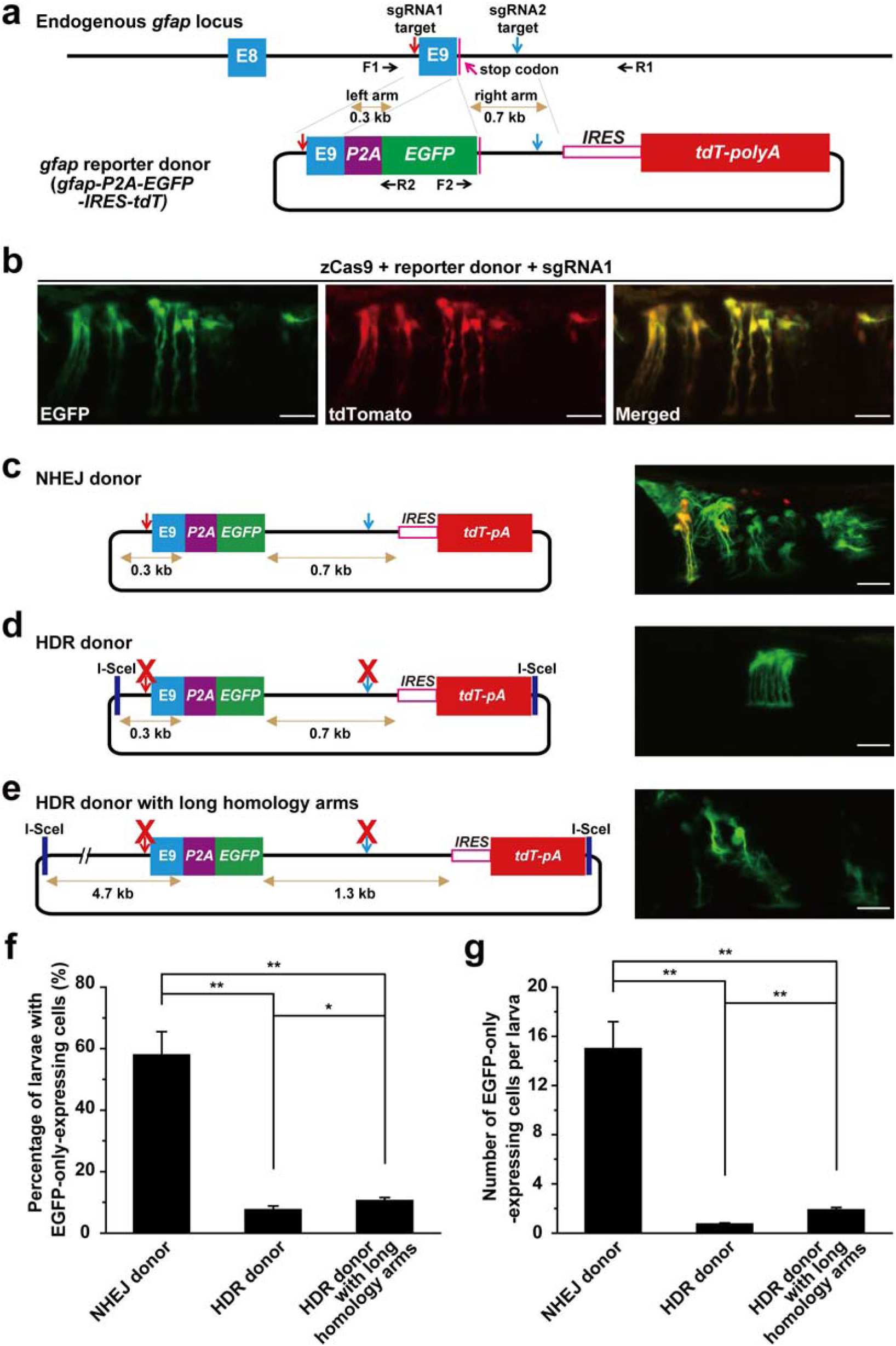
Efficiency comparison of NHEJ- and HDR-mediated DNA replacement at the zebrafish *gfap* locus by using double fluorescence reporter donors. **a** Schematic of the strategy for NHEJ-mediated DNA replacement at the zebrafish *gfap* locus by using a double fluorescence donor vector. The zebrafish *gfap* has 9 exons and E9 represents the 9^th^ exon. The lengths of the left and right homology arms are 324 bps and 702 bps, respectively. IRES: internal ribosome entry site; tdT-polyA: tdTomato connected with a SV40 polyA sequence. **b** Co-expression of EGFP and tdTomato in glial cells when co-injecting the *gfap-P2A-EGFP-IRES-tdT*, zCas9 mRNA and *gfap* sgRNA1. **c** NHEJ-mediated DNA replacement at the *gfap* locus. Left, schematic of the *gfap-P2A-EGFP-IRES-tdT* donor vector for NHEJ-mediated DNA replacement. Right, representative projected *in vivo* confocal image of a 3-dpf larva co-injected with the vector, zCas9 mRNA, *gfap* sgRNA1 and *gfap* sgRNA2. In the vector, the *gfap* sgRNA1 and sgRNA2 target sites are indicated by a red or blue arrow, respectively, and the homology arms are indicated by brown lines with double arrows. **d** HDR-mediated DNA replacement at the *gfap* locus by a donor vector with homology arms which are the same as in **c**. Left, schematic of the HDR-related donor. The PAM of the sgRNA1 and sgRNA2 target sites located in the homology arms of the vector was mutated (red crosses) to prevent CRISPR/Cas9-induced excision. Right, representative projected *in vivo* confocal image of a 3-dpf larva co-injected with the vector, zCas9 mRNA, *gfap* sgRNA1 and *gfap* sgRNA2. (E) HDR-mediated DNA replacement at the *gfap* locus by a donor vector with homology arms longer than those used in **d**. Left, schematic of the HDR-related donor. The lengths of the left and right arms are 4720 bps and 1326 bps, respectively. The PAM of the sgRNA1 and sgRNA2 target sites located in the homology arms of the vector was mutated (red crosses). Right, representative projected *in vivo* confocal image of a 3-dpf larva co-injected with the vector, zCas9 mRNA, *gfap* sgRNA1 and *gfap* sgRNA2. **f, g** Comparison of the efficiencies of NHEJ- and HDR-mediated DNA replacement at the *gfap* locus. **f** Percentage of larvae with EGFP-only-expressing glial cells. **g** Average number of EGFP-only-expressing glial cells per larva. The same data were used in **f, g**. The numbers of larvae examined in “NHEJ donor”, “HDR donor”, and “HDR donor with long homology arms” were 170, 277 and 245, respectively. Scale bars: 20 µm (**b-e**). **\****P* < 0.05; **\*\****P* < 0.01 (unpaired Student’s t-test). Error bars, SEM.

## Discussion

In recent studies, HDR has been used to generate knockin lines of zebrafish with various efficiencies^12-14,24-27^. For most of reported zebrafish lines generated via HDR, single-stranded oligodeoxynucleotide (ssODN) was usually used as a repair template^14,24-27^. This strategy can only achieve genomic alteration with a few of bps such as point mutation and short tag insertion. Long DNA fragment replacement has wide application in gene editing, because it can achieve different types of genomic editing. However, long DNA fragment replacement for zebrafish genome editing is still under development.

Pre-screening via obvious phenotypes or reporter gene expression, both of which imply correct genomic replacement events, was sometimes used to increase screening efficiency in those studies^13,14,24,28^. However, the requirement for pre-screening cannot be met in many cases. Without pre-screening, genomic editing mediated by HDR in zebrafish shows relatively low efficiencies^25-27^, in particular for long DNA fragments^12,14^. This makes it necessary to develop more efficient methods for achieving long DNA fragments replacement.

NHEJ has been recently used to achieve targeted integration in cultured cells, mouse and zebrafish with a high efficiency through inserting a DNA fragment at a single DSB site^21,23,29-33^, but these strategies still cannot be readily applied for sophisticated genomic editing, in particular for conditional knockout. In the present study, by taking advantage of highly active NHEJ-mediated knockin with homology arms and non-coding region targeting, we developed an efficient long DNA fragment replacement method by introducing two DSB sites at non-coding genomic regions via the CRISPR/Cas9 system. With pre-screening, the screen rate of our method for long genomic DNA replacement is 25% (*gfap* reporter allele: 2/8), which is comparable to the reported targeted integrations^13,21,23,28,30,31^. Without pre-screening, the screen rate of our method for long genomic DNA replacement is 9% (*th* conditional allele: 3/41, *kdrl*: 2/25, *tcf3a*: 2/15, *th* reporter allele: 1/11), which is higher than that of HDR-based method in previous studies (4%; *th*: 4/275 and *kcnh6a*: 3/43; see also Refs. 12 and 14). In comparison with HR/HDR-based approaches in a more precise way, we did experiments at the same genomic locus *gfap* and found that the efficiency of NHEJ-mediated DNA replacement is much higher (see Fig. 6f, g). As DSB sites were designed in non-coding regions, NHEJ-induced indels will not shift the reading frame of targeted genes, leading to an increased rate of successful DNA replacement. Interestingly, although NHEJ-mediated DNA repair is independent on homology arms, we found that homology arms extended out of both target sites are important for increasing knockin efficiency (Supplementary Table 2). Based on this data, further studies should test the feasibility of NHEJ-mediated long fragment replacement via universal construct with simply modification by just adding target sites and the short homology arms extended out of target sites. Among our data, only one case is found that the perfect replacement happened without indels at both sides, implying that HDR-mediated replacement cannot be excluded completely by our approach. Thus, the high efficiency and simplicity make our strategy an applicable DNA replacement method for zebrafish and even other organisms.

## Methods

### Zebrafish

Adult zebrafish were maintained with automatic fish housing system (ESEN, China) at 28°C. Embryos were raised under a 14h-10h light-dark cycle in 10% Hank’s solution that consisted of (in mM): 140 NaCl, 5.4 KCl, 0.25 Na2HPO4, 0.44 KH2PO4, 1.3 CaCl2, 1.0 MgSO4 and 4.2 NaHCO3 (pH 7.2). The embryos for imaging experiments were treated with 0.003% 1-phenyl-2-thiourea (PTU, Sigma) from 24 hours post-fertilization to prevent pigment formation. The transgenic line Tg(kdrl:EGFP) and knockin line of Ki(th-P2A-EGFP) were described previously^21,34^, and the transgenic line Tg(dat:mRFP-Cre) was generated by using a modified zebrafish BAC clone which contains the *dat* promoter. To design sgRNAs, AB/WT zebrafish were firstly screened by PCR and sequencing at sgRNA target sites. Zebrafish handling procedures were approved by the Institute of Neuroscience, Chinese Academy of Sciences.

### Production of mRNA and sgRNAs *in vitro*

The zCas9 plasmid pGH-T7-zCas9 was linearized by XbaI and used as a template for Cas9 mRNA *in vitro* synthesis with the mMACHINE T7 Ultra kit (Ambion, USA)^35^. Cre or Flp plasmids (in pcDNA3.1) were linearized by XmaJI and their mRNA were synthesized *in vitro* by mMACHINE T7 Ultra kit. We used the CRISPR/Cas9 design tool provided in the website (http://zifit.partners.org) to select specific targets to minimize off-target effects. The sequences of sgRNA targets tested are listed in Supplementary Table 3. A pair of oligonucleotides containing the targeting sequence of sgRNAs were annealed and cloned at the downstream of the T7 promoter in the pT7-sgRNA vector. The sgRNAs were synthesized by the MAXIscript T7 Kit (Ambion, USA) and were purified by using the mirVana™ miRNA Isolation Kit (Ambion, USA).

### Replacement donor vector construction

Replacement donor plasmids were constructed by standard molecular cloning techniques with restriction enzyme cleavage and DNA ligation. The *th-loxP-exon8-loxP* replacement donor was constructed by ligating three fragments (a left arm, *loxP-exon8-loxP* and a right arm) with the pMD-19T vector via KpnI, BamHI, AgeI and SalI (Takara, Japan). The three fragments were all amplified from the genomic DNA of Ki(th-P2A-EGFP) zebrafish by using the PrimeSTAR HS DNA polymerase (Takara, Japan). The *kdrl-loxP-exon12-frt-SM-frt-loxP* or *tcf3a-loxP-exon4-frt-SM-frt-loxP* replacement donor were constructed by ligating four fragments (a left arm, *loxP-exon12/4, frt-SM-frt*, and a *loxP*-right arm) with the pMD-19T vector via KpnI, SacI, PaeI BglII and SalI. The *th-P2A-EGFP* replacement donor was constructed by modifying the knockin plasmid used in our previous work^21^. The original *P2A-EGFP* fragment was replaced with *P2A-EGFP*-sgRNA2 target by using AflII and AgeI, and then silent mutation of 5 bps in the 3’ side of the *th* CDS was introduced by using the Fast Mutagenesis System (Transgen, China). For constructing different *gfap-P2A-EGFP* replacement donors, the left and right arms of *gfap-P2A-EGFP* were amplified from the genomic DNA of sgRNA target-screened wild-type zebrafish, and the *P2A-EGFP* fragment was obtained from the *th-P2A-EGFP* knockin plasmid. The *gfap-P2A-EGFP-IRES-tdT* donor was constructed by modifying *gfap-P2A-EGFP* plasmid, which ligated the fragment of *IRES-tdT-polyA* with the 3’ side of the shorted right homology arm of the *gfap-P2A-EGFP*. The HDR-related donors were constructed by mutating the PAM regions of *gfap-P2A-EGFP-IRES-tdT* plasmids, and then a pair of head-to-head oriented I-SceI sites were added out of the left arm and *IRES-tdT*.

### Microinjection of one-cell stage embryos

All donor plasmids were purified before microinjection by the Gel Extraction Kit (Qiagen, Germany). Flp mRNA and Cre mRNA were synthesized *in vitro* as zCas9 mRNA. The zCas9 mRNA, sgRNAs, and donor plasmids were co-injected into one-cell zebrafish embryos. Each embryo was injected with 1 nl solution containing 600 ng/μl zCas9 mRNA, 50 ng/μl sgRNA each, and 15 ng/μl donor plasmid into the animal pole. Only 1 U/μl I-SceI enzyme 5 X/μl buffer (NEB) were mixed when co-injecting with the HDR donor. Flp mRNA or Cre mRNA was also injected into one-cell zebrafish embryos with 1 nl containing 100 ng/μl mRNA.

### PCR and sequencing for confirming sgRNA cleavage

Genomic DNA for PCR was extracted from 20 embryos at 3 dpf that were co-injected with zCas9 mRNA and sgRNAs. Fragments containing target sites were amplified by the Takara Ex Taq, and were then cloned by the TA Cloning Kit (Takara, Japan) for sequencing. Generally, we sequenced 20-50 clones for analyzing the cleavage efficiency of each sgRNA.

### Founder screening and the germline mosaicism rate analysis

All F0 zebrafish were screened by out-crossing with AB/WT adults. F1 progenies of Ki(th^EGFP^) or Ki(gfap^EGFP^) founder candidates were screened by EGFP signal at 3 dpf. The ratio of F1 progeny carrying the modification was calculated based on EGFP-positive larvae. 1 - 3 positive larvae were picked up to extract the genomic DNA to determine the integration manner. F1 progenies of Ki(th^fl^-P2A-EGFP) candidates were screened by collecting ∼ 30 F1 progenies of each candidate for genomic DNA extraction and PCR identification. The ratio of F1 progeny carrying the modification was analyzed by PCR analysis of 48 F1 progenies one by one. F1 progenies of Ki(kdrl^fl^) or Ki(tcf3a^fl^) candidates were screened first by DsRed signal in heart, then 1-3 positive larvae were picked up for genome extraction and PCR identification. The ratio of F1 progeny carrying the modification was calculated based on DsRed-positive larvae. The genomic DNA of positive F1 larvae was also used for confirmation of the integration manner.

### Verification of NHEJ-mediated DNA replacement

The genomic DNA of 1 - 20 3-dpf F1 embryos of each F0 were extracted. The fragment of 5’ and 3’ junctions were amplified respectively with the primer pair of F1/R2 and F2/R1, whereas the full length product of the entire edited region was amplified with F1/R1 (Supplementary Table 4). The size of all PCR products was first examined by gel electrophoresis, and the full length products amplified by F1/R1 were sequenced directly or cloned and sequenced. The sequencing data were used to analyze the integration manner at junction sites or the excision at *frt*/*loxP* sites.

### In vivo confocal imaging

For *in vivo* confocal imaging, larvae at 3 or 5 dpf were immobilized in 1.2 % low melting-point agarose without paralysis or anesthetics. Imaging experiments were performed under a 20X water-immersion objective by using a Fluoview 1000 two-photon microscope (Olympus, Japan)^36^. The spatial resolution of all images was 1024 × 1024 pixels.

### Statistics

Statistical analysis was performed using unpaired Student’s *t*-test. The *P* value less than 0.05 were considered to be statistically significant. All results are represented as mean ± SEM.

## Acknowledgements

This work was supported by the star program of Shanghai Jiao Tong University (YG2019ZDA20), Key Research Program of Frontier Sciences (QYZDY-SSW-SMC028), Strategic Priority Research Program (XDB32010200) and International Partnership Program (153D31KYSB20170059) of Chinese Academy of Sciences, and Shanghai Municipal Science and Technology Major Project (18JC1410100, 2018SHZDZX05).

## Author contributions

J.L.D. and S.Y.G. conceived the project. S.Y.G., L. Jiang, C.H. and J.L.D. designed experiments. S.Y.G., Y.G.R. and J. Li built donor constructs. S.Y.G., J.W.B. and J.B.C carried out PCR diagnosis. S.Y.G. and J. Li performed all other experiments and analysis. S.Y.G. and J.L.D. wrote the paper with inputs from L. Jiang and C.H.

## Competing interests

The authors declare no competing interests.

