## Supplemental information for "Efficient replacement of long DNA fragments via non-homologous end joining at non-doding regions"

Gu *et al.*

Supplementary Information

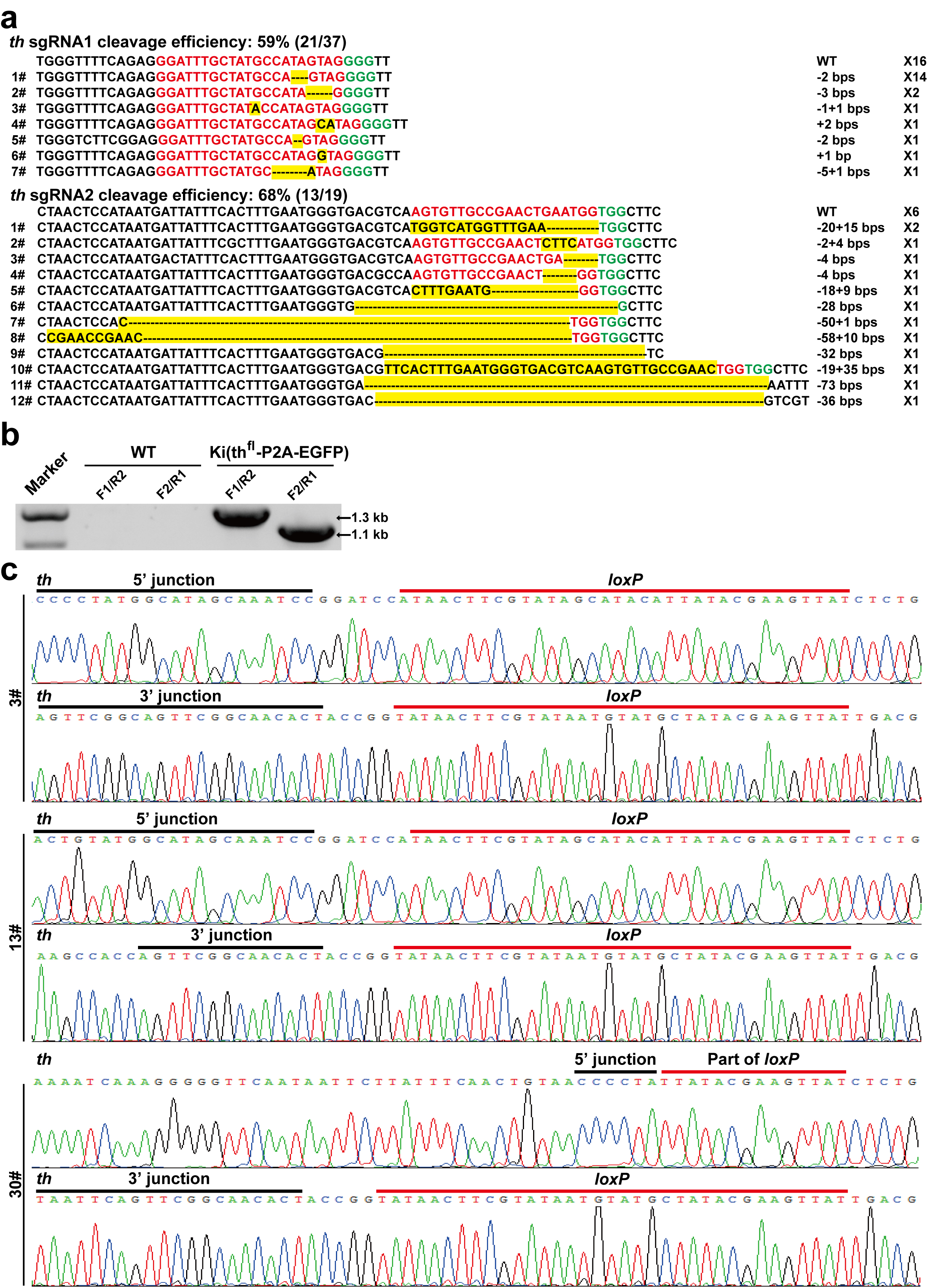

**Supplementary Figure 1. Cleavage efficiency of *th* sgRNAs and verification of *th* conditional alleles.** **a,** Cleavage efficiency of the sgRNA1 and sgRNA2 targeting the zebrafish *th* locus. The indel mutations are highlighted in yellow, and the PAM and sgRNA target sequences are shown in green and red, respectively. The numbers of insertion (+) and/or deletion (-) of base pairs (bps) are shown at the right. **b,** PCR analysis of the 5’ and 3’junctions of the targeted *th* locus. The primers of F1, R1, F2, and R2 are shown in **Fig. 1a**. A 1.3-kb band was amplified by using the 5’ junction primers F1 and R2, and a 1.1-kb band was amplified by using the 3’ junction primers F2 and R1. **c,** Sequencing reports of the entire edited region PCR product at the 5’ and 3’ junction sites of the progenies of three Ki(th^fl^-P2A-EGFP) founders.

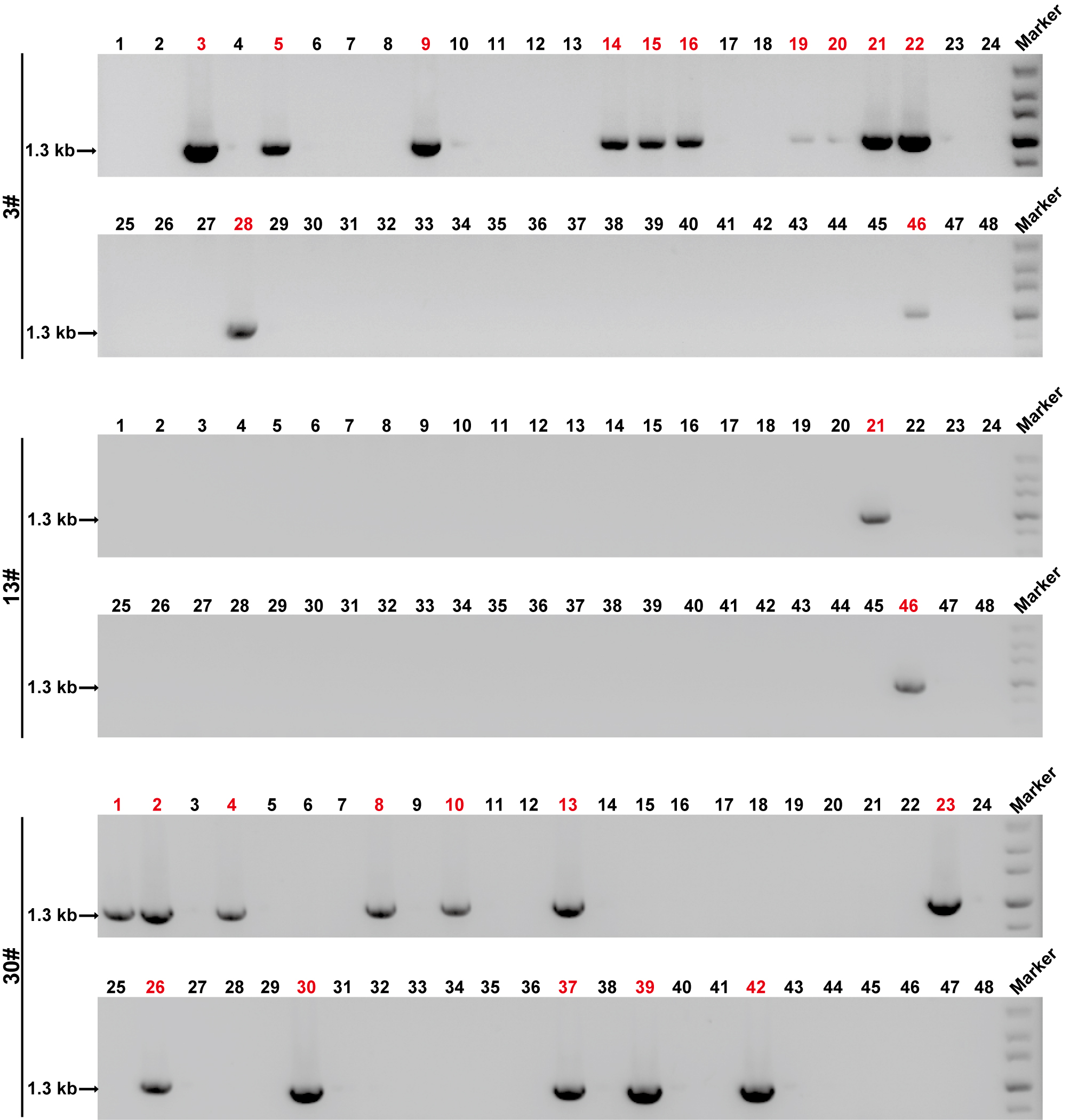

**Supplementary Figure 2. Germline mosaicism rate analysis of the three Ki(th^fl^-P2A-EGFP) founders by PCR.** The 5’ junction primers F1 and R2 were used to amplify DNA fragments from 48 individual F1 of each founder. The number in red indicates that the corresponding progeny carries a conditional allele.

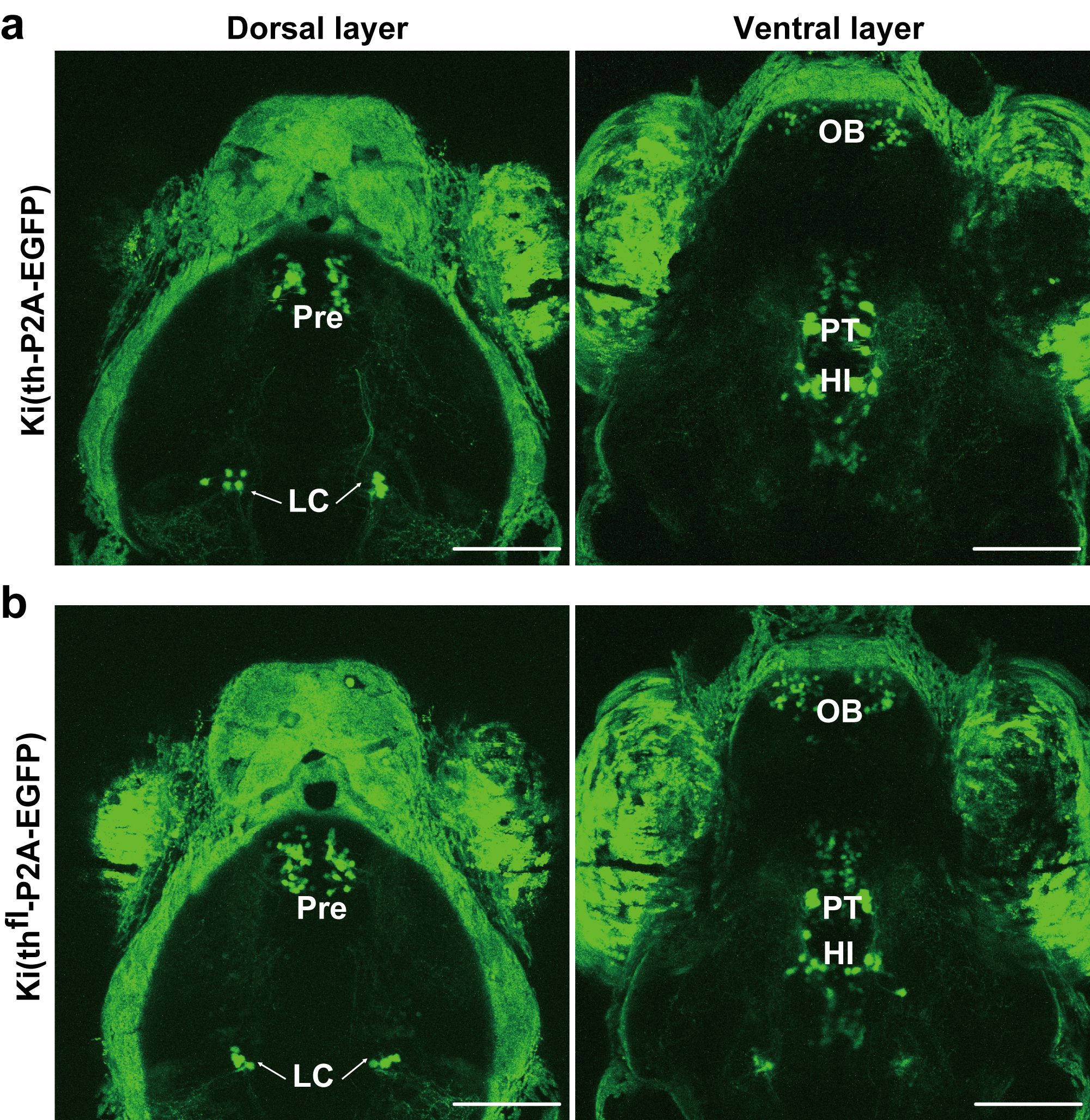

**Supplementary Figure 3. Similarity of EGFP expression patterns between the larvae of Ki(th-P2A-EGFP) and Ki(th^fl^-P2A-EGFP).** **a,** Representative projected *in vivo* confocal images (dorsal view) of a 3-dpf Ki(th-P2A-EGFP) larva at the dorsal and ventral layers. **b,** Representative projected *in vivo* confocal images (dorsal view) of a 3-dpf Ki(th^fl^-P2A-EGFP) larva at the dorsal and ventral layers. HI, intermediate hypothalamus; LC, locus coeruleus; OB, olfactory bulb; Pre, pretectum; PT, posterior tubercular. Scale bars: 100 µm (**a**, **b**).

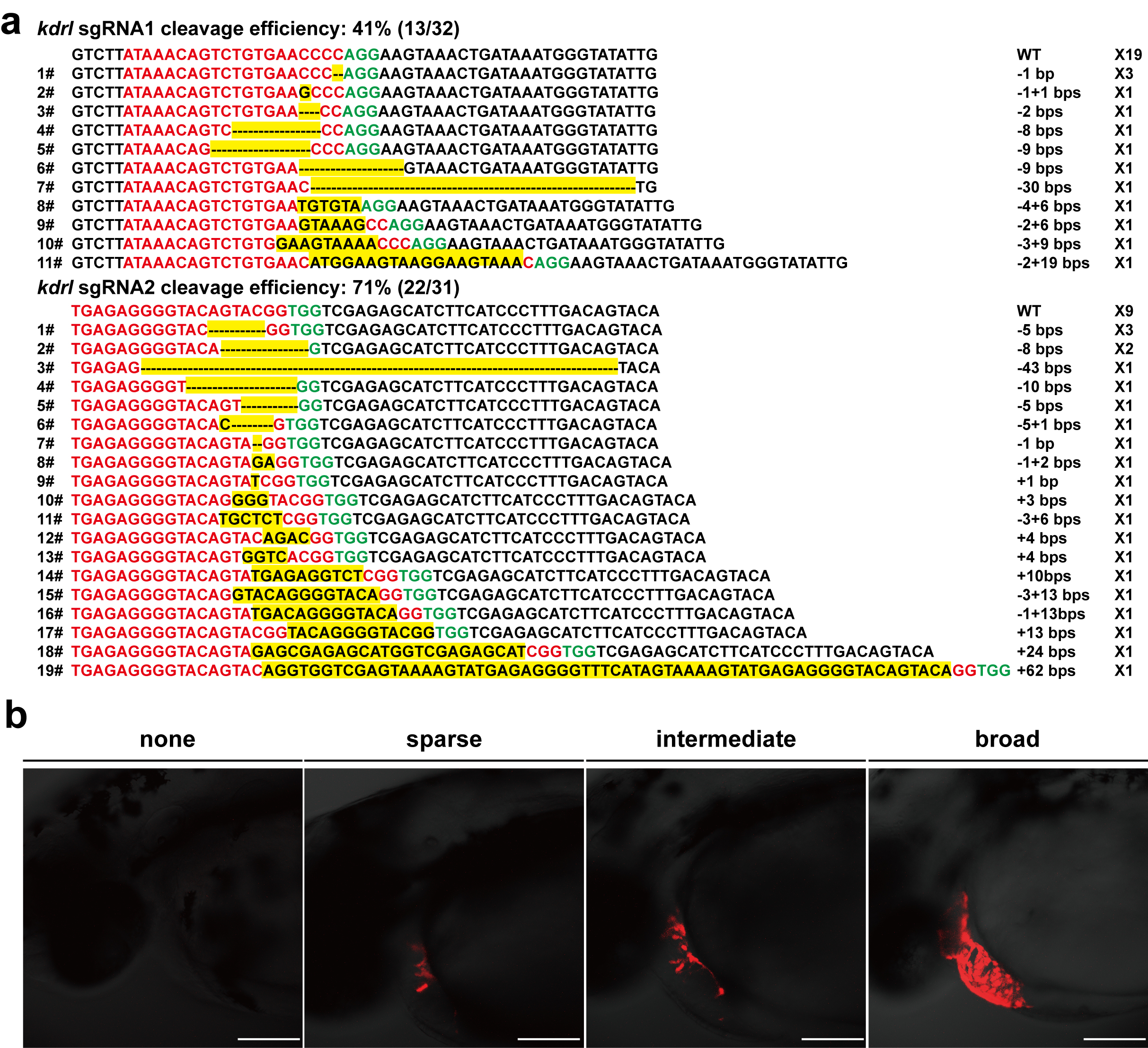

**Supplementary Figure 4. Cleavage efficiency of *kdrl* sgRNAs and transient expression of the selective marker.** **a,** Cleavage efficiency of the sgRNA1 and sgRNA2 targeting the zebrafish *kdrl* locus. The indel mutations are highlighted in yellow, and the PAM and sgRNA target sequences are shown in green and red, respectively. The numbers of insertion (+) and/or deletion (-) of bps are shown at the right. **b,** Differential expression of SM in the heart of F0 larvae. Scale bars: 100 µm.

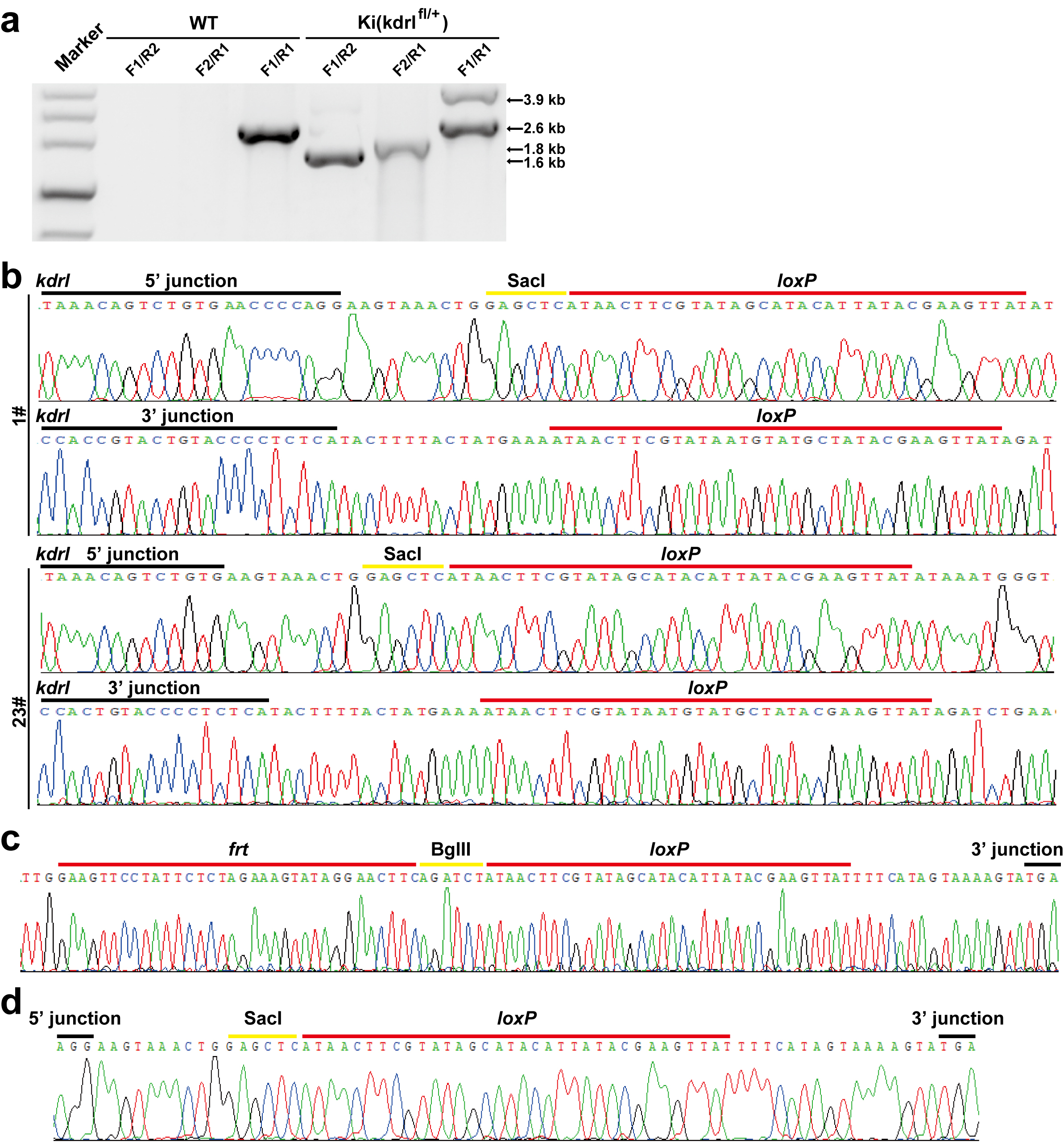

**Supplementary Figure 5. Verification of *kdrl* conditional alleles. a,** The entire edited region PCR analysis of the 5’ and 3’junctions of the targeted *kdrl* locus. The primers of F1, R1, F2, and R2 are shown in Figure 2. A 1.6-kb band was amplified by using the 5’ junction primers F1 and R2, a 1.8-kb band was amplified by using the 3’ junction primers F2 and R1, and a 2.6-kb band and a 3.9-kb band were amplified by using the primers F1 and R1. **b,** Sequencing reports of the entire edited region PCR product at the 5’ and 3’ junction sites of the progenies of two Ki(kdrl^fl^) founders (1# and 23#). **c,** Sequencing report showing of the entire edited region PCR product SM deletion by Flp. **d,** Sequencing report of the additional 2.1-kb product showed in Figure 2D, confirming the correct deletion by Cre.

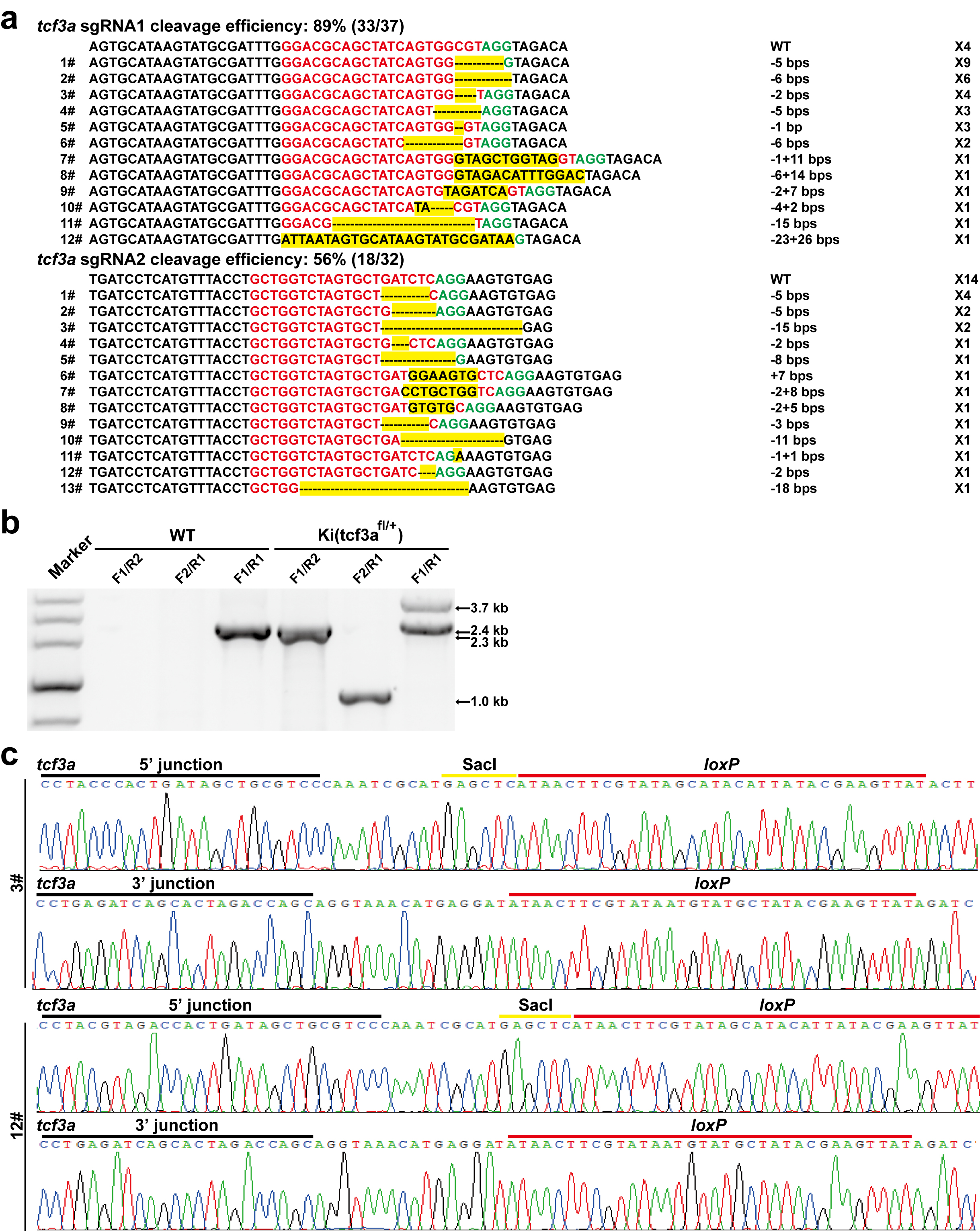

**Supplementary Figure 6. Cleavage efficiency of *tcf3a* sgRNAs and verification of *tcf3a* conditional alleles.** **a,** Cleavage efficiency of the sgRNA1 and sgRNA2 targeting the zebrafish *tcf3a* locus. **b,** The entire edited region PCR analysis of the 5’ and 3’junctions of the targeted *tcf3a* locus. **c,** Sequencing reports of the entire edited region PCR product at the 5’ and 3’ junction sites of the progenies of two Ki(tcf3a^fl^) founders (3# and 12#).

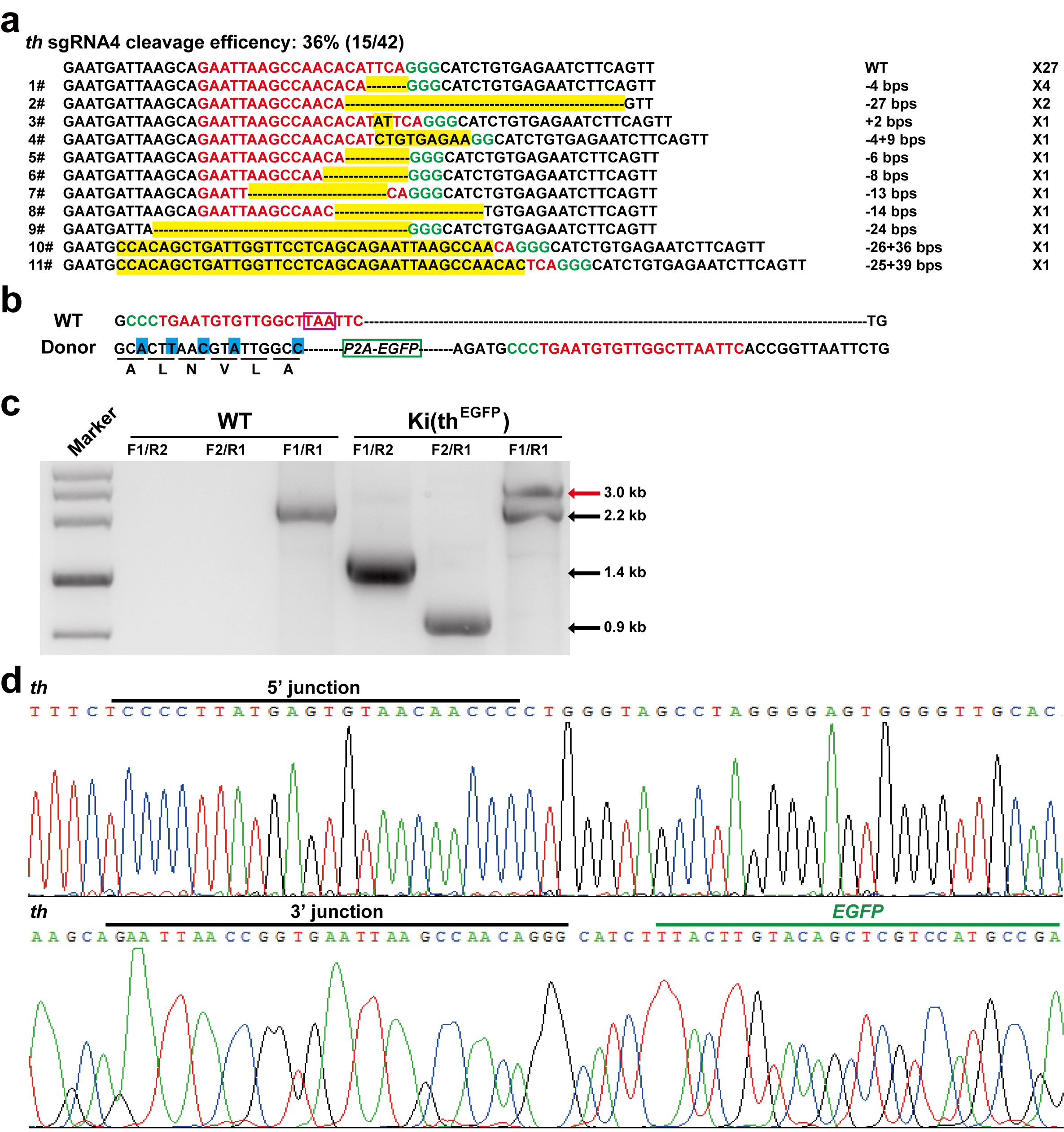

**Supplementary Figure 7. Cleavage efficiency of *th* sgRNA and confirmation of the Ki(th^EGFP^) reporter line. a,** Cleavage efficiency of the sgRNA4 targeting the zebrafish *th* locus. The indel mutations are highlighted in yellow, and the PAM and sgRNA target sequences are shown in green and red, respectively. The numbers of insertion (+) and/or deletion (-) of base pairs are shown at the right. **b,** sgRNA4 target sequences in the *th* locus and the reporter donor. Silent mutations are highlighted in blue. The *th* stop codon and *P2A-EGFP* are indicated by a pink or green box, respectively. Please note, the sgRNA3 with a cleavage efficiency of 83% was the same as that used in Ref. 21. **c,** The entire edited region PCR verification of DNA replacement at the *th* locus by using the primers (F1, R1, F2 and R2) shown in **Fig. 4a**. For the endogenous *th* locus, a 2.2-kb band was amplified by using F1 and R1. For a targeted *th* locus, 1.4-kb band was amplified by using the 5’ junction primers F1 and R2, a 0.9-kb band was amplified by using the 3’ junction primers F2 and R1, and a 3.0-kb band (red arrow) was amplified by using F1 and R1. F1 progenies of a Ki(th^EGFP^) founder (#4) were used. **d,** Sequencing reports of the entire edited region PCR product at the 5’ and 3’ junction sites of the progeny of the Ki(th^EGFP^) founder.

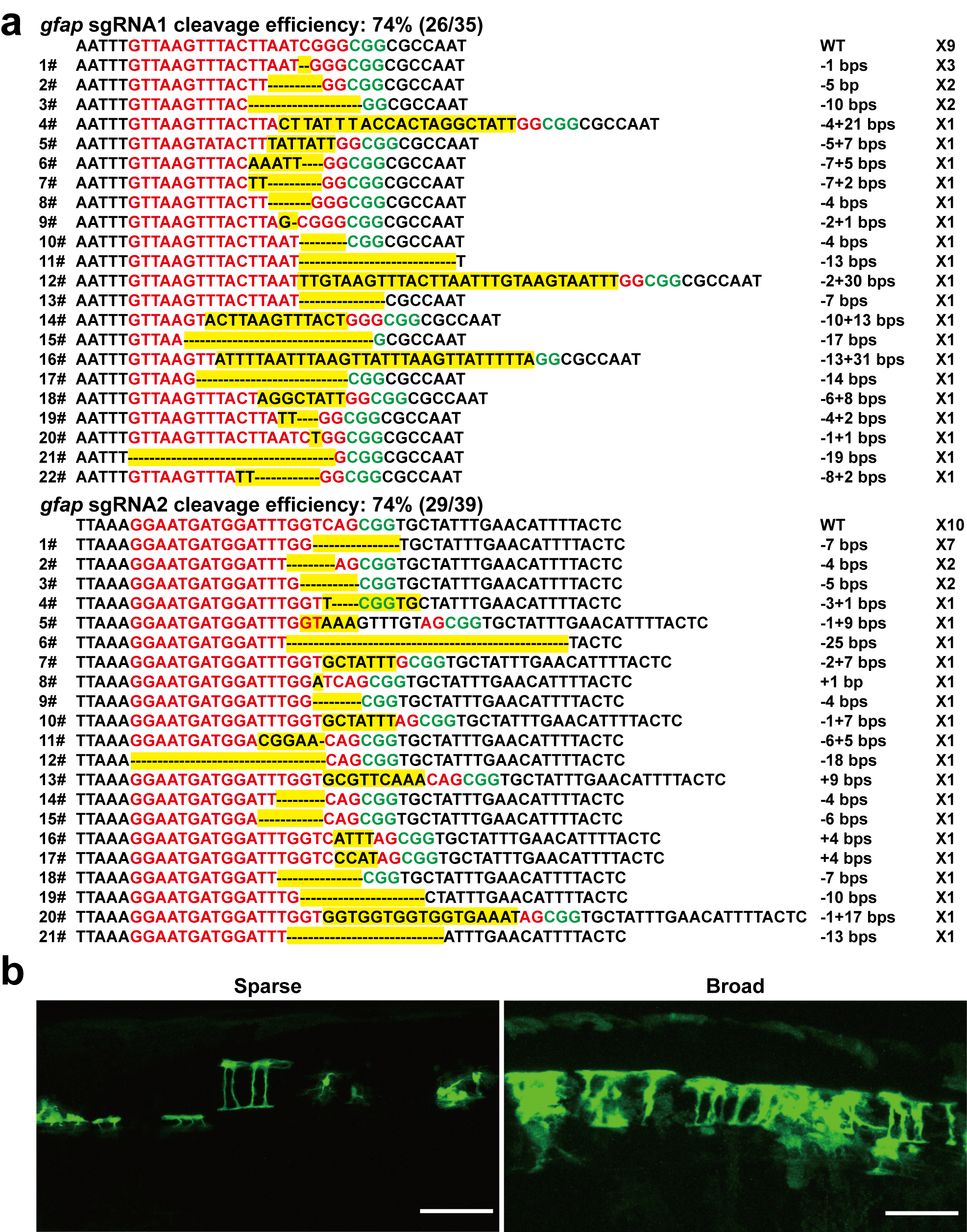

**Supplementary Figure 8. Cleavage efficiency of *gfap* sgRNA and verification by transient expression in F0 larvae.** **a,** Cleavage efficiency of the sgRNA1 and sgRNA2 targeting the zebrafish *gfap* locus. **b,** Transient EGFP expression in glial cells when co-injecting the *gfap-P2A-EGFP*, zCas9 mRNA, and sgRNA1 and sgRNA2 of *gfap*. Scale bars: 50 µm.

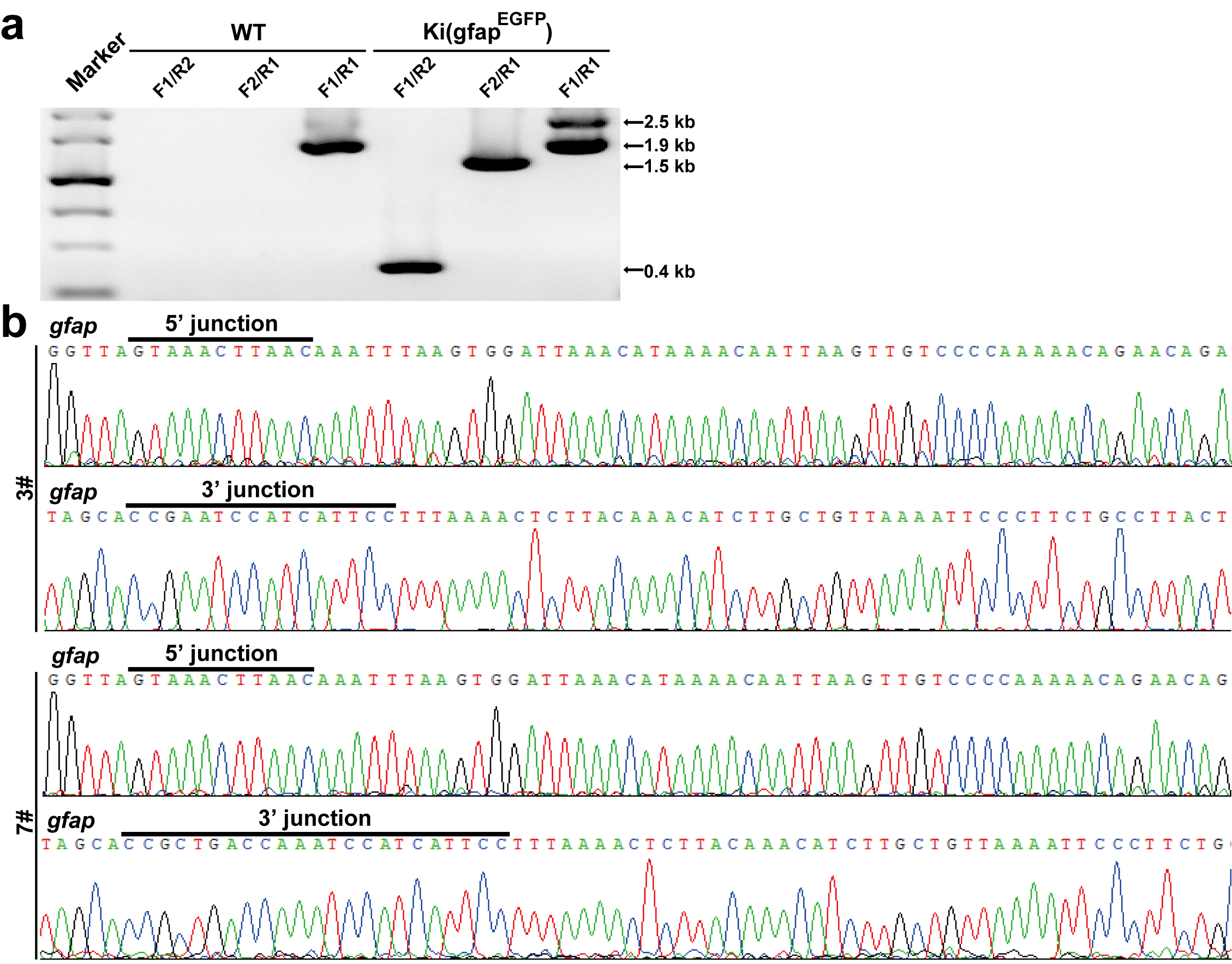

**Supplementary Figure 9. Confirmation of the Ki(gfap^EGFP^) reporter line. a,** The entire edited region PCR verification of DNA replacement at the *gfap* locus. **b,** Sequencing reports of the entire edited region PCR product at the 5’ and 3’ junction sites of the progenies of two Ki(gfap^EGFP^) founders (3# and 7#).

**Supplementary Table 1.** **Founder-screening rate of different edited alleles by NHEJ-mediated DNA replacement.**

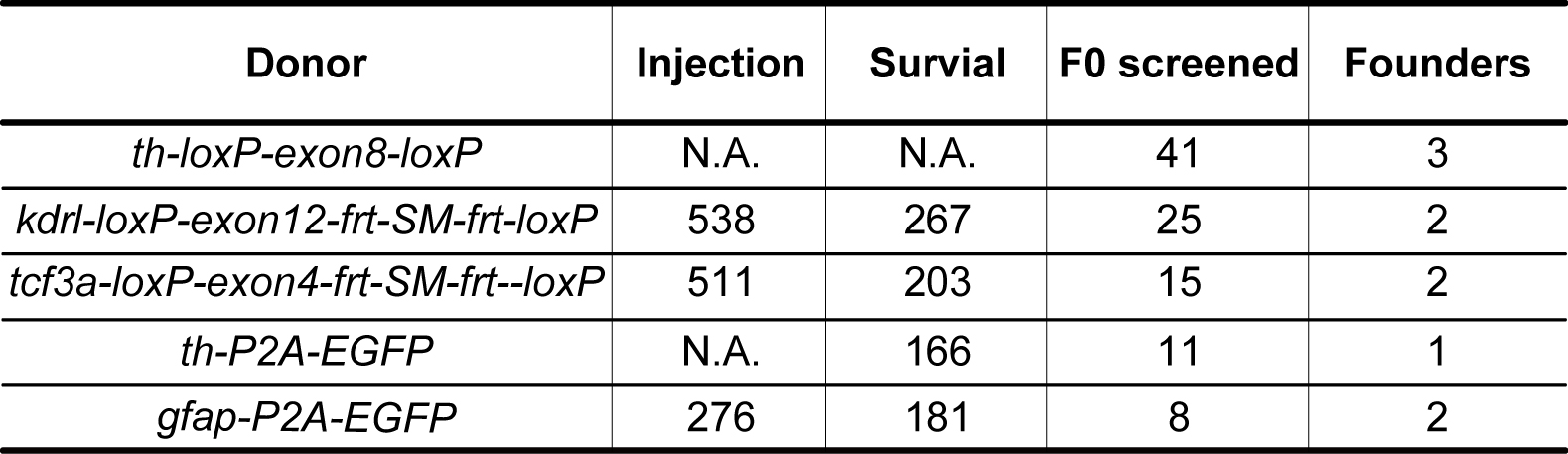

N.A.: not available.

**Supplementary Table 2. Efficiency of knockin with different lengths of homology arms at the *gfap* locus.**

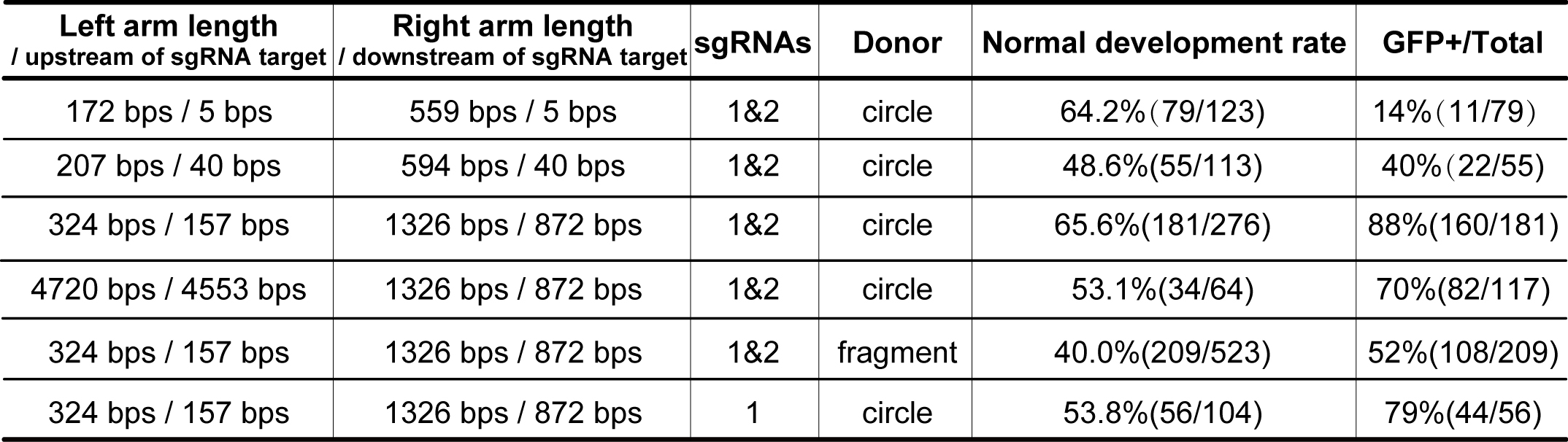

**Supplementary Table 3.** **Target sequences of all tested sgRNAs.**

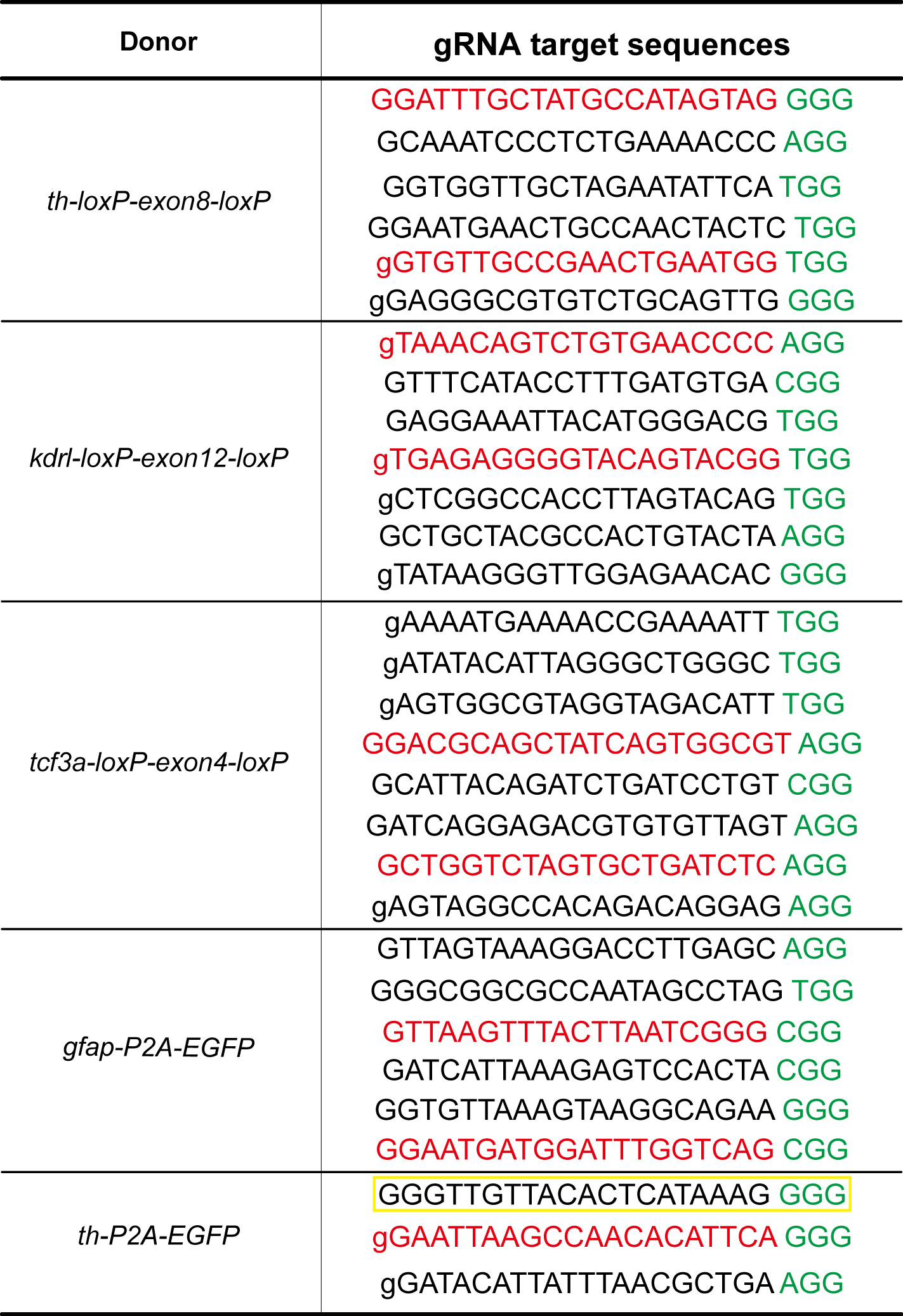

The sequences in red were used to generate DNA replacement-mediated knockin zebrafish in this study, and yellow box represents an efficient target sequence which was used in a previous study (Ref. 21). The green characters represent PAM sequences. The "g" was added for synthesizing sgRNAs.

**Supplementary Table 4. Sequences of primers used.**

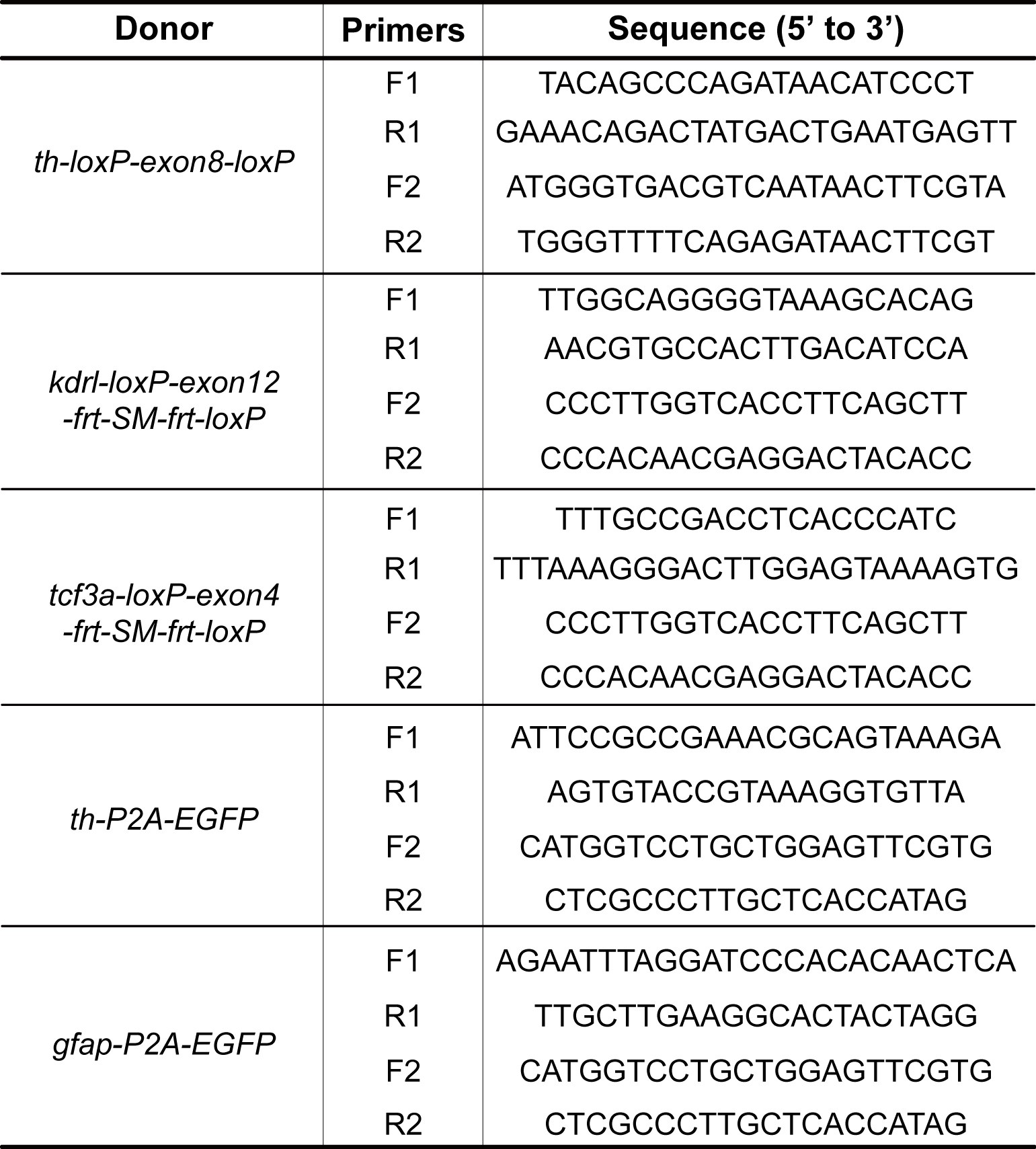
